# Circulating apoB-containing lipoproteins constituted blood-borne innate immune barrier limits ZIKV dissemination and congenital infection risk

**DOI:** 10.64898/2026.09.15.751642

**Authors:** Juncheng Chen, Dawei Lv, Wenlu Che, Yujie Chen, Yuanfei Zhu, Na Liu, Nihua Dong, Luhe Gan, Xinya Ji, Jiayu Qian, Guanglei Zhou, Yiming Tong, Jin Zhong, Mingbing Xiao, Ralf Bartenschlager, Gang Long

## Abstract

Following establishment of primary viremia, flaviviruses require prompt penetration through vascular endothelium to achieve systemic dissemination, yet most infections remain asymptomatic or self-limited. Canonical defenses, including complement and phagocyte clearance, cannot fully explain this containment, suggesting an additional blood-borne antiviral barrier. Here, we identify apoB-containing lipoproteins as potent restriction factors against Zika virus (ZIKV), a devastating danger of global pregnancy. By displaying surface-accessible phosphatidylserine (PS), they competitively occupy host PS receptors and block entry by viruses exploiting apoptotic mimicry, including flaviviruses, Ebola, and chikungunya virus. In vivo, increasing apoB-lipoproteins enhances serum antiviral activity and limits systemic ZIKV dissemination. This barrier extends to the maternal–fetal interface, where elevated maternal LDL attenuates vertical transmission and fetal abnormalities. Its efficacy is modulated by gestational fluctuations in maternal LDL level and low abundance in fetal circulating LDL. Collectively, apoB-containing lipoproteins form a constitutive blood-borne innate immune barrier that restrains apoptotic-mimicry viruses during the intravascular phase, limits viral dissemination and reduces congenital infection risk. This study provides a rationale for developing diet augmentation strategy to strengthen this innate barrier in at-risk individuals.

## INTRODUCTION

Blood-borne dissemination is a defining step in many severe viral infections. After breaching anatomic barriers, pathogenic viruses can enter the circulation and gain access to distant organs, enabling systemic disease, neuroinvasion, vascular leakage or congenital infection. This route is particularly consequential for re-emerging arthropod-mediated and blood-borne flaviviruses, which cause recurrent outbreaks and a broad spectrum of severe disease. Dengue virus (DENV) causes hundreds of millions of infections each year and can progress to severe plasma leakage, shock and death^1,2^, whereas West Nile virus (WNV) and tick-borne encephalitis virus (TBEV), can cause neuroinvasive disease^3,4^. During pregnancy, Zika virus (ZIKV) can reach placental and fetal tissues and cause fetal growth restriction, microcephaly, and other congenital abnormalities^5,6^. Thus, systemic infection requires circulating virions to remain infectious, evade innate clearance, and access susceptible tissues.

An emerging entry strategy that supports this dissemination is apoptotic mimicry. By exposing phosphatidylserine (PS) on the virion surface, flaviviruses engage host PS-recognition pathways, including TIM and TAM family receptors^7–9^. Through this strategy, virions exploit machinery used for uptake of dying cells, thereby promoting attachment, internalization and immune evasion. Remarkably, the broad distribution of PS-recognition receptors on endothelial and epithelial cell populations expand flaviviral tropism and promote vascular escape during systemic dissemination^7,10,11^.

Following inoculation by blood-feeding arthropods, flaviviruses replicate locally before entering the circulation during primary viremia^12^. In blood, canonical blood-borne defenses including complement, natural antibodies, and mononuclear phagocytes contribute to virion clearance^13,14^. Therefore prompt escape from intravascular space is essential for flaviviruses to seed peripheral tissues and cause severe diseases. Clinically, most flavivirus infections remain asymptomatic or self-limited^12,15,16^, suggesting that circulating virions are restrained in blood. Therefore, rather than serving merely as a conduit for viral dissemination, blood constitutes an intrinsic and potent barrier poised to intercept virus seeding and corresponding pathological outcome.

The majority of ZIKV infections are asymptomatic, with less than 0.03% incidence of ZIKV-associated Guillain-Barre syndrome. In contrast, maternal-fetal transmission may occur in all trimesters of pregnancy in 30% of fetuses from ZIKV infected mothers. Approximately 35% infection positive fetuses showed fetal loss and congenital Zika syndrome including microcephaly^17^. ZIKV infection still poses a devastating danger of global pregnancy. Here, using ZIKV and animal models, we identified circulating apoB-containing lipoproteins as abundant endogenous fluid-phase restriction factors against ZIKV infection. By competing for PS-recognition receptors, this lipoprotein barrier restricts viral entry at the vascular interface and reduces systemic and vertical ZIKV dissemination. These findings expand the concept of blood-borne innate immunity. Declining of apoB-containing lipoproteins during the first trimester is a neglected determinant of congenital ZIKV infection risk.

## RESULTS

### Serum apoB-containing lipoproteins restrict ZIKV infection

Systemic viral dissemination requires viremia and productive peripheral infection before viral seeding of target tissues^18,19^. We first asked whether serum contains intrinsic antiviral activity against ZIKV. To avoid interference from FBS-derived components in routine virus preparations, we generated serum-free ZIKV stocks (**Fig. 1A**). These stocks were then incubated with heat-inactivated sera from uninfected mammalian hosts, including FBS, mouse serum (MS) and human serum (HS). Even in the absence of active complement, all three sera showed dose-dependent suppression of ZIKV infection (**Fig. 1B**). Human serum displayed the highest antiviral activity, achieving maximal blockade of viral infection at physiological concentrations. This inhibition was corroborated across multiple quantitative assays, including reduced intracellular viral RNA loads (**Fig. 1C**), decreased infection rates (**Fig. 1D, E**), and diminished production of infectious progeny (**Fig. 1F, G**). Similar inhibition was observed in A549 and Vero E6 cells, where serum addition during infection significantly reduced intracellular viral genome copies in both cell lines (**Fig. S1A, B**).

**Fig. 1.**
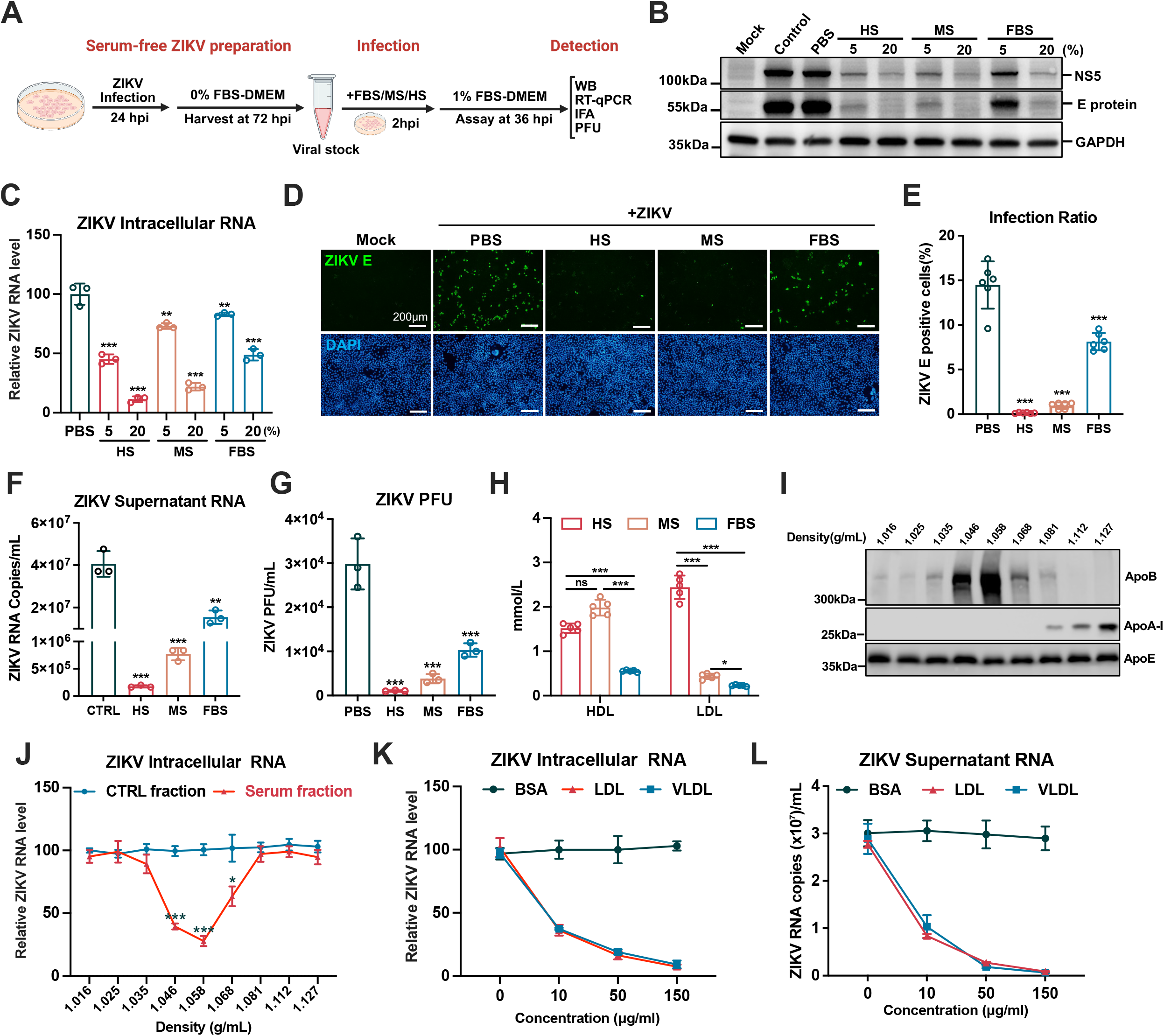
Identification of circulating lipoproteins as intrinsic restriction factors of ZIKV infection. **A**, Schematic diagram of the experimental workflow. Serum-free ZIKV stocks were generated by propagating passage-1 (P1) virions in Vero E6 cells, with medium replaced by serum-free DMEM at 24 hours post-infection (hpi). Viral supernatant was harvested at 48 hpi. For infection assays, Huh7 cells were co-incubated with ZIKV and indicated sera for 2 h, washed, and maintained in 1% DMEM until 36 hpi. **B–G**, Huh7 cells were infected with serum-free ZIKV (MOI = 0.1) in the presence of indicated concentrations of heat-inactivated fetal bovine serum (FBS), pooled mouse serum (MS), or pooled healthy human serum (HS). Assays were performed at 36 hpi. **B**, Immunoblot analysis of intracellular ZIKV NS5 and envelope (E) proteins. **C**, Intracellular ZIKV RNA levels normalized to *GAPDH*, measured by Q-RT-qPCR. **D**, Representative immunofluorescence images of ZIKV E-positive cells. **E**, Quantification of the infection rate (percentage of ZIKV-positive cells) across six randomly selected fields using ImageJ. **F**, Extracellular viral RNA levels in supernatant. **G**, Infectious progeny viral titers. **H**, HDL and LDL cholesterol concentrations in FBS, MS, and HS, determined via colorimetric quantification kits. **I**, Immunoblot analysis of apolipoprotein B (APOB), APOA1, and APOE across human serum fractions separated by density gradient ultracentrifugation. **J**, Intracellular ZIKV RNA levels in Huh7 cells treated with 20% (v/v) of the indicated serum fractions from **I** at 36 hpi. **K**, **L**, Intracellular (**K**) and extracellular (**L**) ZIKV RNA levels in Huh7 cells supplemented with the indicated concentrations of bovine serum albumin (BSA) or purified human VLDL or LDL at 36 hpi. Data are presented as means ± s.d. from three independent infection experiments. Statistical significance was determined using one-way ANOVA with Dunnett’s multiple comparisons test (**C**, **E**–**G**, **J-L**) or Tukey’s multiple comparisons test (**H**). \**P* < 0.05; \*\**P* < 0.01; \*\*\**P* < 0.001. The schematic diagram in **A** was created with BioRender.com.

The FBS, MS and HS used above were obtained from ZIKV-naive hosts, suggesting that this activity was mediated by an intrinsic serum component. Given the correlation between serum anti-ZIKV activities and low-density lipoprotein (LDL) levels among human serum, murine serum and fetal bovine serum (**Fig. 1H**), we hypothesized that these metabolic carriers might drive the observed antiviral activity. We therefore fractionated human serum using density gradient ultracentrifugation. Immunoblotting for APOB and APOAI confirmed separation of lipoprotein subclasses (**Fig. 1I**). Subsequent functional assessment revealed that fractions enriched with LDL and VLDL (density 1.016–1.068 g/mL) exhibited the strongest suppression of ZIKV infection (**Fig. 1J**). Consistent with the fractionation results, purified LDL and VLDL significantly and dose-dependently suppressed ZIKV infection as measured by intracellular ZIKV RNA levels (**Fig. 1K**) and progeny viral titers (**Fig. 1L**). In addition, LDL and VLDL also inhibited ZIKV infection in other cell lines (**Fig. S1C-F**). To further validate the antiviral contribution of endogenous apoB-containing lipoproteins in serum, we partially depleted apoB-positive particles from human serum using anti-apoB antibody. ApoB depletion reduced serum ApoB abundance and attenuated serum-mediated restriction of ZIKV infection (**Fig. S2A, B**). Notably, LDL and VLDL blocked ZIKV infection at 150 μg/mL, corresponding to approximately 0.39 mmol/L cholesterol equivalent. This concentration is substantially below the circulating LDL range typically maintained in human blood^20^, indicating that physiological apoB-lipoprotein abundance is sufficient to provide a high-capacity antiviral barrier within vascular compartment.

### LDL and VLDL restrict TIM-1-dependent ZIKV attachment and entry

To define the stage of the viral life cycle targeted by LDL and VLDL, we performed time-of-addition assays (**Fig. 2A**). LDL and VLDL demonstrated significant antiviral effects at entry stage, whereas addition after viral internalization failed to restrict infection. Progeny viral titers were similarly reduced (**Fig. 2B**). Using a 4 °C virus binding assay, we confirmed that lipoprotein treatment significantly reduced surface-bound ZIKV RNA on Huh7 cells (**Fig. 2C**). To exclude effects on post-entry replication, we constructed a ZIKV subgenomic replicon system with or without a polymerase-deficient mutation (ZIKV-SGR, ZIKV-SGR ΔGDD)^21,22^ (**Fig. S3A**). Lipoprotein treatment had no impact on viral RNA replication (**Fig. S3B**), and genome translation demonstrated by comparable luciferase activities (**Fig. S3C**).

**Fig. 2.**
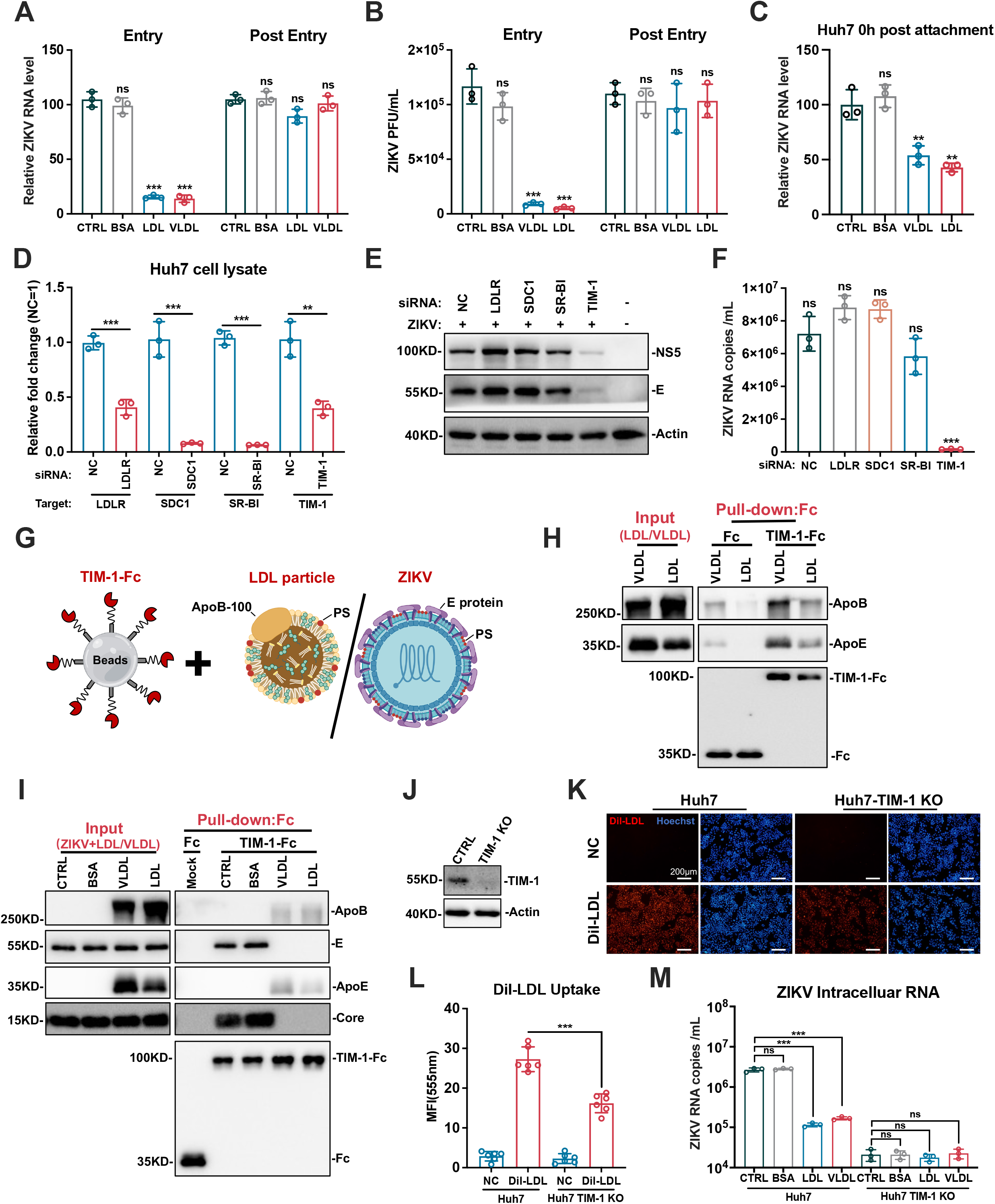
LDL and VLDL restrict ZIKV cellular entry by competitively engaging the TIM-1 receptor. **A**, **B**, Huh7 cells were infected with ZIKV (MOI = 0.1) and supplemented with PBS or 150 μg/mL BSA, VLDL, or LDL either during the initial viral incubation (Entry) or after removing the viral inoculum (Post-entry). At 36 hpi, intracellular ZIKV RNA levels normalized to *GAPDH* were quantified by Q-RT–qPCR (**A**), and progeny viral titers were determined by plaque assay (**B**). **C**, Viral attachment assay. Huh7 cells were pre-chilled to 4 °C and co-incubated with ZIKV (MOI = 10) and 150 μg/mL of the indicated proteins. Relative RNA levels of surface-bound ZIKV were quantified by RT–qPCR. **D–F**, Huh7 cells were transfected with a non-targeting control siRNA (NC) or siRNAs targeting the indicated genes. Assays were performed at 36 hpi. Shown are the relative mRNA knockdown efficiencies of target genes (**D**), ZIKV NS5 and E protein expression via immunoblotting (**E**), and relative intracellular ZIKV RNA levels (**F**). **G**, Schematic of the *in vitro* co-incubation and pull-down assay. **H**, **I**, Protein G magnetic beads coated with 10 μg of recombinant human TIM-1-Fc were used to capture 20 μg of purified VLDL or LDL (**H**), or 5×10^6^ PFU of ZIKV in the presence of 20 μg of the indicated lipoproteins (**I**). Captured complexes were analyzed by immunoblotting for APOB, APOE, ZIKV E, and core proteins. **J**, Immunoblot validation of TIM-1 knockout (KO) efficiency in a monoclonal Huh7 cell line. **K**, **L**, Representative fluorescence images (**K**) and corresponding quantification (**L**) of DiI-LDL internalization in wild-type (WT) and TIM-1-KO Huh7 cells. Cells were incubated with 5 μg/mL DiI-LDL for 2 h. Mean fluorescence intensity (**L**) was quantified across six randomly selected fields of view per condition. **M**, WT and TIM-1-KO Huh7 cells were infected with ZIKV (MOI = 0.1) in the presence of 150 μg/mL BSA, VLDL, or LDL. Intracellular ZIKV RNA levels were determined by Q-RT–qPCR at 36 hpi. Data are presented as means ± s.d. from three independent infection experiments. Statistical significance was determined using two-way ANOVA with Šídák’s multiple-comparison test for two-factor designs (**A**, **B**, **M**), one-way ANOVA with Dunnett’s multiple comparisons test (**C, D**, **F**), or two-tailed unpaired Student’s t-test (**L**). \**P* < 0.01; \*\*\**P* < 0.001; ns, not significant. The schematic diagram in **G** was created with BioRender.com.

We next investigated the molecular mechanism of this entry restriction. ZIKV particles immunoprecipitated with neutralizing antibodies (Z23) did not co-purify with LDL or VLDL, ruling out direct virion binding and infectivity neutralization (**Fig. S4A, B**). This suggested that either lipoproteins masked essential host factor required for ZIKV entry or ZIKV utilized host factors that mediate LDL uptake. Therefore, a siRNA screen targeting LDLR, Syndecan1, SR-BI and T-cell immunoglobulin and mucin domain 1 (TIM-1) was performed (**Fig. 2D**), only TIM-1 knockdown reduced ZIKV infection (**Fig. 2E, F**). Using recombinant TIM-1-Fc fusion proteins, we captured significant amounts of APOB and APOE from LDL or VLDL, indicating direct binding capability between TIM-1 and apoB-containing lipoprotein particles (**Fig. 2G, H**). TIM-1-Fc efficiently captured ZIKV viral structural proteins, whereas the addition of physiological concentrations of LDL or VLDL competitively displaced the virus from the receptor (**Fig. 2I**). TIM-1 knockout Huh7 cells exhibited a marked reduction in fluorescent DiI-LDL internalization compared to wild-type cells, indicating that TIM-1 can mediate LDL uptake (**Fig. 2J-L)**. In TIM-1 knockout Huh7 cells, the antiviral protective effect of LDL was completely abrogated (**Fig. 2M**). Together, these findings establish TIM-1 as a shared PS-recognition receptor exploited by ZIKV, LDL and VLDL, thereby enabling these lipoprotein to competitively restrict TIM-1-dependent ZIKV entry.

### LDL and VLDL engage PS receptors through surface-exposed PS

To elucidate the molecular determinant underlying LDL-mediated antiviral activity, we focused on phosphatidylserine (PS), the canonical ligand for TIM-1. This receptor recognizes PS through a conserved metal-ion-dependent binding pocket^23,24^. To test whether this interface is required for ZIKV attachment, we constructed a PS-binding-deficient mutant (TIM-1-WF/AA)^25^. Loss of PS recognition eliminated TIM-1 binding to ZIKV virions (**Fig. 3A**), and competitive blockade with PS liposomes significantly reduced ZIKV infection (**Fig. 3B**). Wild-type TIM-1 efficiently captured APOB and APOE from LDL and VLDL, whereas TIM-1-WF/AA failed to bind these targets (**Fig. 3C**). To assess functional uptake, we expressed these specific receptors in 293T cells (**Fig. 3D**). Wild-type TIM-1 enhanced DiI-LDL internalization, but neither the TIM-1-WF/AA mutant nor the PS-independent receptor DC-SIGN facilitated LDL uptake (**Fig. 3E, Fig. S5A**). PS liposomes also inhibited TIM-1-mediated LDL entry (**Fig. 3F, Fig. S5B**). Similarly, masking of surface PS on LDL with Annexin V reduced TIM-1-mediated uptake by approximately 50% (**Fig. S5C, D**) and Annexin V treatment also suppressed ZIKV infection (**Fig. S5E**). These results indicate that surface-exposed PS allows apoB-containing lipoproteins to engage TIM-1 and compete with ZIKV for the same receptor interface.

**Fig. 3.**
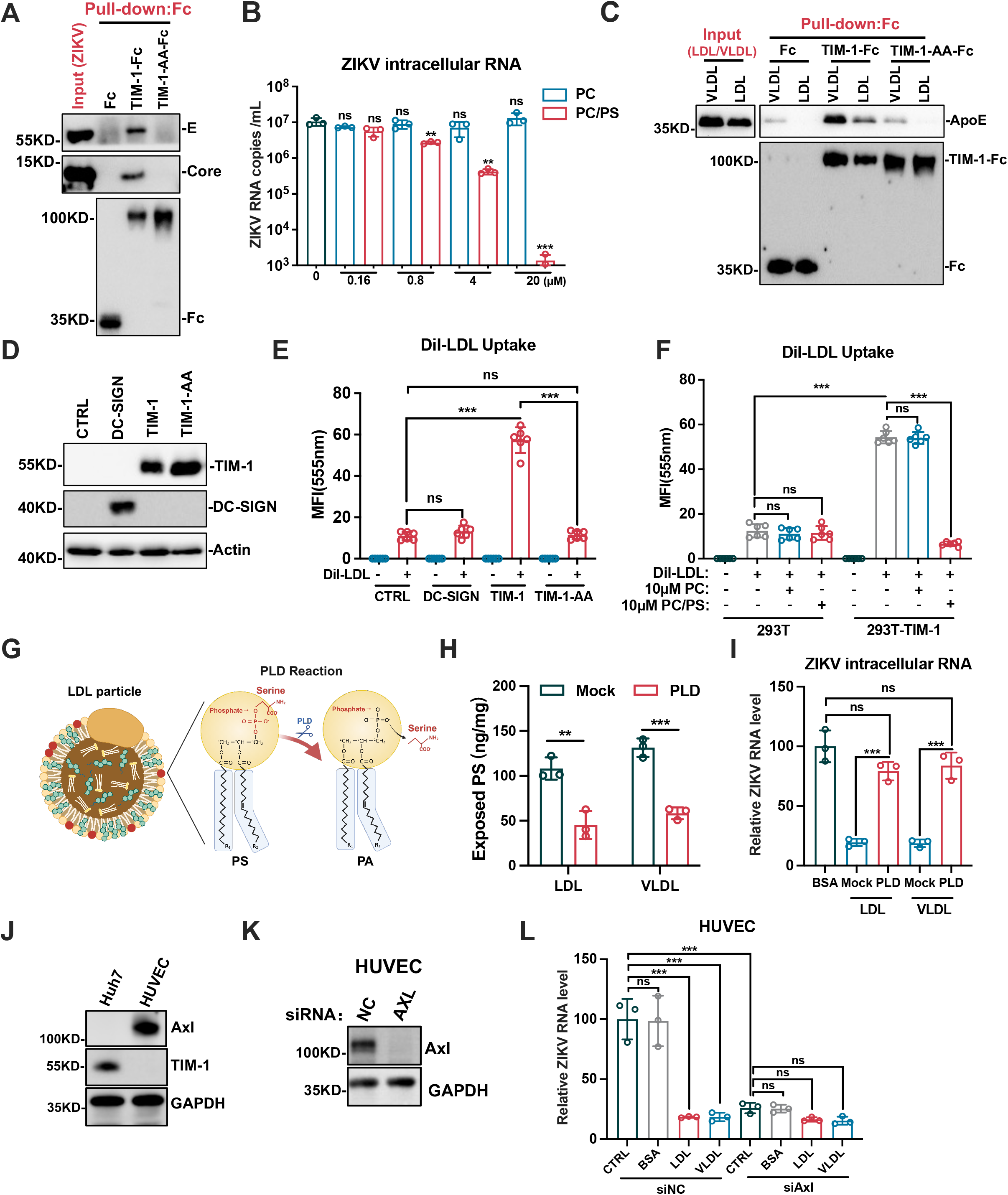
The lipoprotein decoy mechanism relies on surface-exposed phosphatidylserine to restrict ZIKV entry. **A**, *In vitro* viral pull-down assay. Protein G magnetic beads coated with 10 μg of recombinant human IgG1-Fc, TIM-1-Fc, or the PS-binding-deficient TIM-1-WF/AA-Fc mutant were utilized to capture 5 × 10^6^ PFU of ZIKV, followed by immunoblot analysis for ZIKV E and core proteins. **B**, Huh7 cells were infected with ZIKV in the presence of PC or PC/PS liposomes. Relative intracellular ZIKV RNA levels normalized to *GAPDH* were determined by Q-RT–qPCR at 36 hpi. **C**, Immunoblot analysis of APOB and APOE from 20 μg of VLDL or LDL captured by the indicated Fc protein-coated beads. **D**, Immunoblot analysis of TIM-1 and DC-SIGN expression in the indicated 293T cell lines. **E**, Quantification of DiI-LDL mean fluorescence intensity across six randomly selected fields in the indicated 293T cell lines following a 2-h incubation. **F**, DiI-LDL internalization (mean fluorescence intensity) in wild-type and TIM-1-overexpressing 293T cells in the presence of 20 μM PC or PC/PS liposomes. **G**, Schematic representation of the phospholipase D (PLD) enzymatic reaction. **H**, Surface PS levels on VLDL and LDL (50 μg) treated with water (mock) or 100 U of PLD, quantified via PS specific ELISA. **I**, Huh7 cells were infected in the presence of mock- or PLD-treated lipoproteins. Relative intracellular ZIKV RNA levels were measured at 24 hpi. J. Immunoblot analysis of TIM-1 and AXL expression in Huh7 cells and HUVECs. K, Immunoblot analysis of AXL knockdown efficiency in HUVECs transfected with non-targeting control siRNA (NC) or AXL-specific siRNA (AXL). **L,** Relative intracellular ZIKV RNA levels in HUVECs (siNC vs. siAXL). Cells were infected (MOI = 0.1) in the presence of indicated concentrations of bovine serum albumin (BSA), VLDL or LDL.Data are presented as means ± s.d from three independent experiments. Statistical significance was determined using one-way ANOVA with Dunnett’s multiple comparisons test (**B**), two-way ANOVA with Šídák’s multiple-comparison test for two-factor experiments (**E**, **F**, **H**, **I**), or one-way ANOVA with Tukey’s multiple comparisons test (**L**). \**P* < 0.05; \*\**P* < 0.01; \*\*\**P* < 0.001; ns, not significant. The schematic diagram in **G** was created with BioRender.com.

Vascular endothelial cells form a major interface between circulating virions and peripheral tissues^26^. We next examined this effect in human umbilical vein endothelial cells (HUVECs)^27,28^. Unlike Huh7 cells, which predominantly expressed TIM-1, HUVECs showed robust AXL expression with negligible TIM-1 (**Fig. 3G**). LDL and VLDL significantly reduced ZIKV infection in HUVECs, and AXL depletion impaired ZIKV infection and attenuated LDL/VLDL-mediated restriction (**Fig. 3H, I**), indicating that apoB-containing lipoproteins restrict endothelial infection through an AXL-dependent PS-entry pathway. These findings suggest that apoB-containing lipoproteins broadly antagonize PS receptors and this lipoprotein-mediated restriction prevents hematogenous dissemination.

### LDL and VLDL specifically restrict viruses exploiting apoptotic mimicry

In light of the highly conserved nature of apoptotic mimicry as a viral entry strategy^7,10,29^, we investigated whether lipoprotein-mediated receptor antagonism confers broad-spectrum antiviral restriction. Here, we included multiple members of the Flaviviridae family (DENV-2, WNV, TBEV)^7^, a picornavirus (HAV)^30^, a filovirus surrogate (Ebola VLPs)^31^, and an alphavirus (CHIKV)^32^. To evaluate the mechanistic specificity of this decoy effect, we performed parallel challenge assays using viruses (SARS-CoV-2 and HSV)^33–35^ that employ PS-independent entry pathways. Consistent with our previous finding on ZIKV, supplementation with apoB-containing lipoproteins inhibited infection of DENV, WNV, TBEV, HAV,Ebola and CHIKV (**Fig. 4A**-**F**). In contrast, they did not affect SARS-CoV-2 or HSV-1 infection **(Fig. 4G, H)**. Taken together, these data show that LDL and VLDL specifically restrict viruses that rely on PS recognition pathways for cell entry, but not viruses that use alternative entry strategies.

**Fig. 4.**
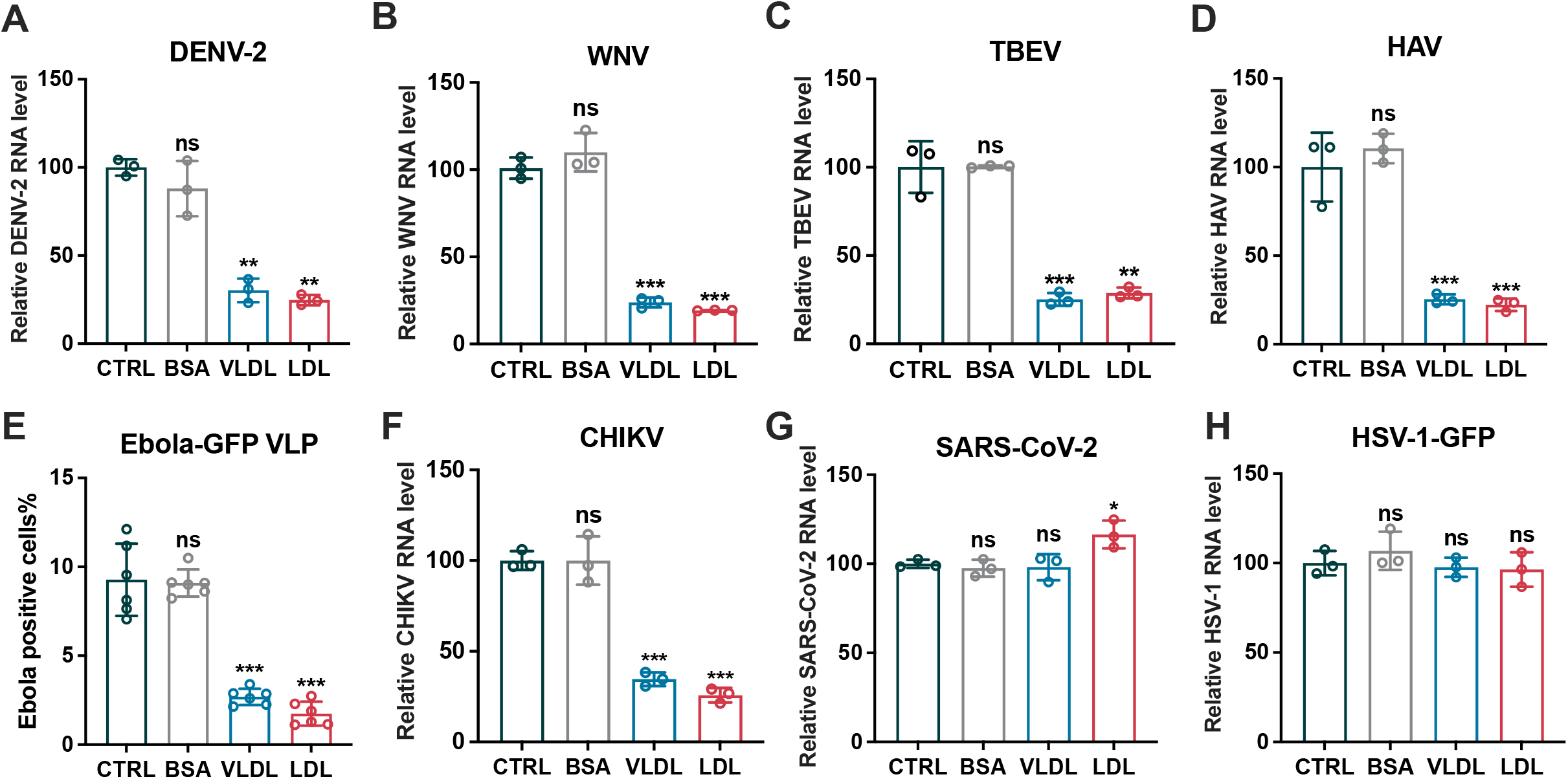
Lipoprotein-mediated restriction is specific to viruses exploiting apoptotic mimicry. **A**–**D**, Huh7 cells were infected with DENV-2 (**A**), WNV (**B**), TBEV (**C**), or HAV (**D**) at an MOI of 0.1 in the presence of 150 μg/mL BSA, VLDL, or LDL. Relative intracellular viral RNA levels normalized to *GAPDH* were quantified by Q-RT–qPCR in at 36 hpi. **E**, Infection rate, defined as the percentage of GFP-positive cells (quantified via ImageJ), in Huh7-4PX cells inoculated with GFP-reporter Ebola VLPs in the presence of the indicated proteins at 48 hpi. **F**, Relative intracellular CHIKV RNA levels normalized to *GAPDH* in Huh7 cells at 24 hpi. **G**, Relative intracellular SARS-CoV-2 RNA levels in Vero E6 cells at 24 hpi. **H**, Relative intracellular HSV-1 mRNA levels normalized to *GAPDH* in Huh7 cells at 36 hpi. Data are presented as means ± s.d. from three independent experiments. Statistical significance was determined using one-way ANOVA with Dunnett’s multiple comparisons test (**A**–**H**). \**P* < 0.05; \*\**P* < 0.01; \*\*\**P* < 0.001; ns, not significant.

### Elevation of apoB-containing lipoprotein improves serum anti-ZIKV capability

To establish the physiological relevance of this LDL-driven decoy mechanism, using multiple hyperlipidemia models, we next investigated how natural fluctuations in circulating lipoproteins impact serum anti-ZIKV capability. First, we examined *Ldlr*^−/-^mice^36,37^, which showed markedly elevated serum LDL levels (**Fig. 5A**) and increased serum anti-ZIKV activity compared with WT mice (**Fig. 5B**). Density-gradient fractionation of WT and *Ldlr*^−/-^ sera further showed that antiviral activity was concentrated in LDL-enriched fractions and increased with LDL abundance (**Fig. 5C, D)**. A similar phenotype was observed in *Apoe*^−/-^ mice, with increased LDL abundance accompanied by enhanced serum-mediated restriction of ZIKV (**Fig. 5E, F**). We further examined a diet-induced obesity (DIO) model, in which elevated serum LDL was similarly associated with stronger anti-ZIKV activity (**Fig. 5G**, **H**). At last, LDL abundance of human serum positively correlated with the ex vivo capacity to restrict ZIKV infection (**Fig. 5E**). Taken together, these results established that circulating LDL level is the major determinant of serum antiviral efficacy.

**Fig. 5.**
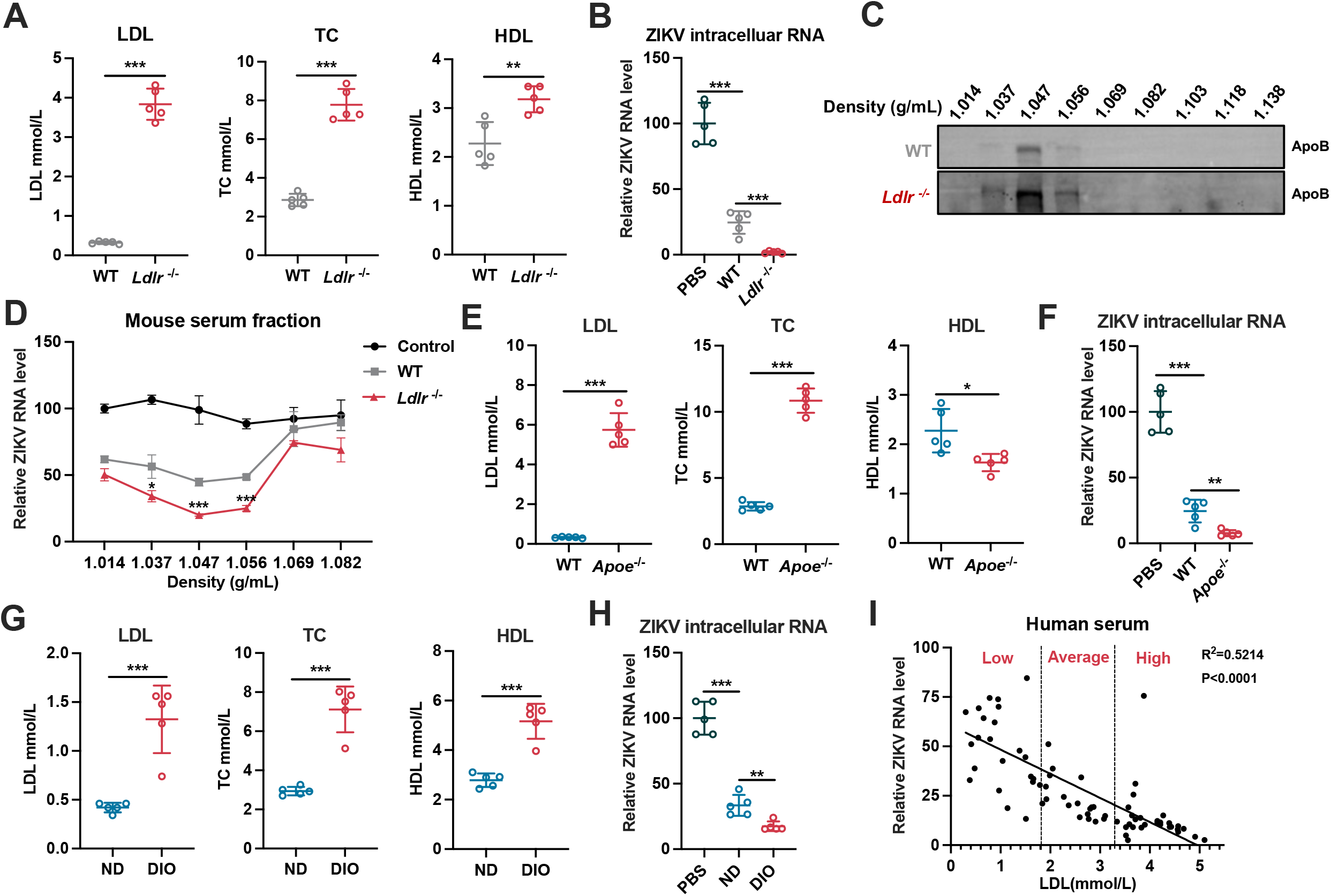
Serum lipoprotein concentrations positively correlate with ex vivo antiviral efficacy. **A**, Serum total cholesterol (TC), HDL, and LDL concentrations in wild-type (WT) and *Ldlr^−/-^* male mice (n = 5). **B**, Huh7 cells were infected in the presence of 20% indicated murine sera in **A** (n = 5). Relative intracellular ZIKV RNA levels at 36 hpi. were quantified. **C**, Immunoblot analysis of APOB across mouse serum fractions separated by density gradient ultracentrifugation. **D**, Huh7 cells were infected in the presence of 20% indicated serum fractions from **C.** Relative intracellular ZIKV RNA levels at 36 hpi were quantified **E,** Serum TC, HDL, and LDL in 8-week-old WT and Apoe-/- mice (n = 5). **F,** Huh7 cells were infected in the presence of 20% indicated murine sera in E(n = 5). Relative intracellular ZIKV RNA levels at 36 hpi. **G,** Serum TC, HDL, and LDL concentrations in 16-week-old normal diet (ND) control and diet-induced obese (DIO) mice (n = 5), determined using an automated rodent biochemical analyzer. **H,** Huh7 cells were infected in the presence of 20% indicated murine sera in G (n = 5). Relative intracellular ZIKV RNA levels at 36 hpi. Data are means ± s.d. from three independent experiments. Statistical significance was determined using two-tailed unpaired Student’s t-tests (**A–D**). **I**, Huh7 cells were infected in the presence of 10% heat-inactivated clinical human serum(n = 73 individuals) versus baseline serum LDL concentrations.Relative intracellular ZIKV RNA levels at 36 hpi. Simple linear regression of relative intracellular ZIKV RNA levels was demonstrated (I). Statistical significance was determined using two-tailed unpaired Student’s t-tests (**A–H**) or simple linear regression with Pearson correlation (**I**), \**P* < 0.05; \*\**P* < 0.01; \*\*\**P* < 0.001.

### Elevated circulating LDL intercept systemic ZIKV dissemination

Given that the *Ldlr^−/-^* model provides a more specific expansion of the circulating LDL fraction, we selected this genetic background to evaluate the functional role of LDL in restricting ZIKV dissemination in vivo. We generated *Ldlr*^−/-^ models on an AG129 background^18,19^ (**Fig. S6A-C**), resulting in *Ldlr*^−/-^ AG129 mice. AG129, *Ldlr*^−/-^AG129 and *Ldlr*^−/-^ AG129 with high-cholesterol diet (HCD) models were included for in vivo ZIKV challenge, recapitulating low, average and high LDL levels in human sera (**Fig. 6A)**. Following ZIKV infection, *Ldlr*^−/-^ AG129 mice exhibited delayed weight loss and reduced mortality. Additionally, *Ldlr*^−/-^ AG129 mice with HCD supplementation demonstrated further protection (**Fig. 6B, C**). Serum lipid profile showed that ZIKV infection inherently doubled basal LDL levels; within this infected context, *Ldlr*^−/-^ mice displayed a 10-fold increase in serum LDL, compared to wild-type controls, whereas HCD feeding drove a 30-fold elevation (**Fig. 6D**). ZIKV RNA burdens in the serum, spleen, kidney, brain, and liver decreased in a dose-dependent manner corresponding to serum LDL levels (**Fig. 6E**). Histopathological examination confirmed that ZIKV-induced severe neuronal shrinkage and hyperchromasia were significantly mitigated in mice with elevated LDL (**Fig. 6F, G**). Collectively, these findings established circulating LDL as a blood-borne antiviral barrier that limits systemic ZIKV dissemination.

**Fig. 6.**
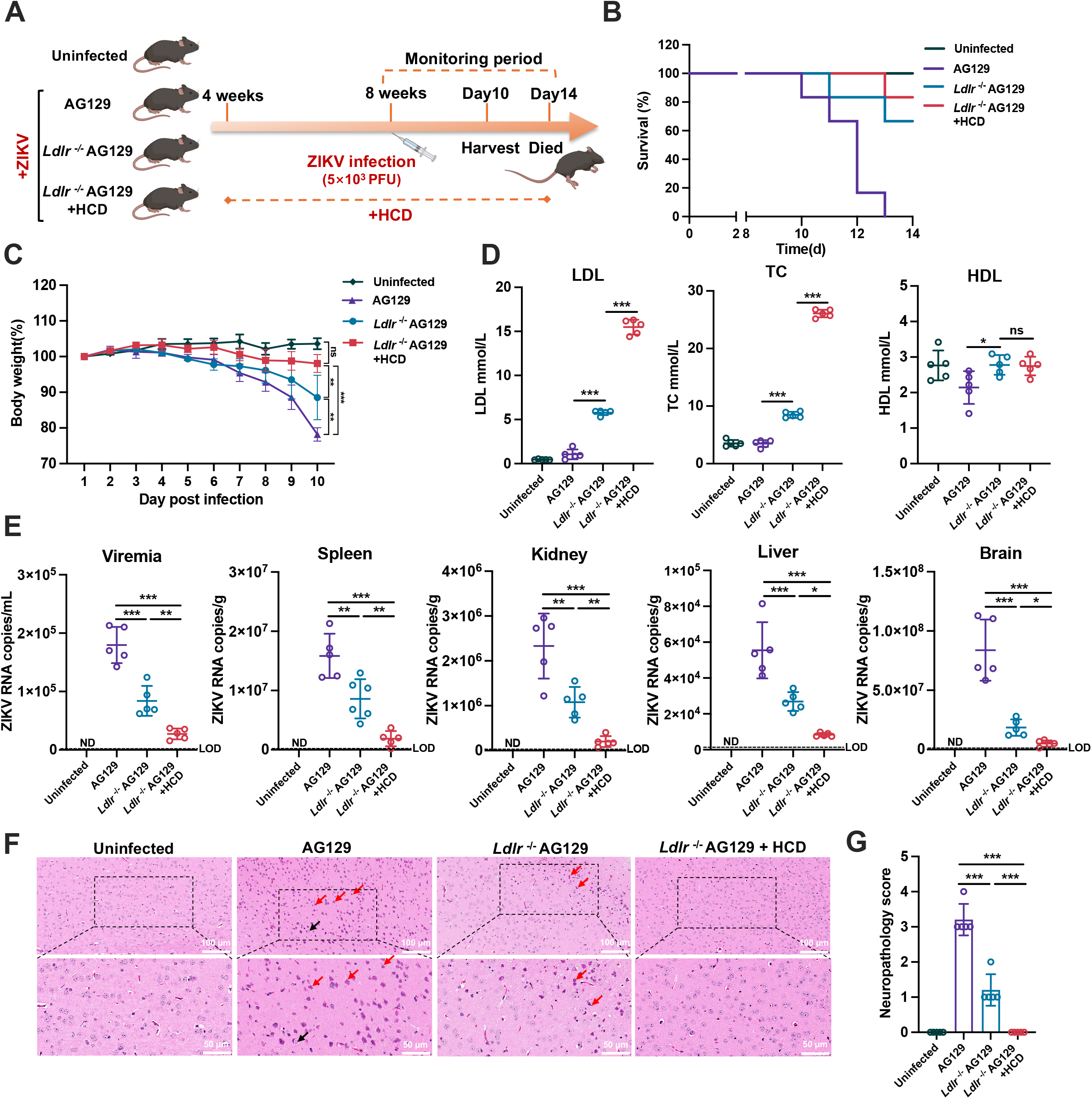
Elevated circulating LDL intercepts systemic ZIKV infection in vivo. **A**, Schematic diagram of *in vivo* experimental design. Male mice on an AG129 immunodeficient background were challenged subcutaneously (s.c.) with 5×10^3^ PFU of ZIKV. **B**, **C**, Kaplan–Meier survival curves (**B**, n = 6) and body weight changes (**C**, n = 5) of the indicated mouse cohorts following ZIKV infection. **D**, Serum TC, HDL, and LDL concentrations across all experimental groups at 10 days post-infection (dpi) (n = 5). **E**, Viral loads in target tissues (copies per gram) and serum (copies per milliliter) quantified by Q-RT–qPCR at 10 dpi (n = 5 per group). **F**, **G**, Representative hematoxylin and eosin (H&E)-stained brain sections at 10 dpi (**F**), showing abundant shrunken and hyperchromatic neurons with reduced cell volumes and indistinct nucleocytoplasmic borders (red arrows), alongside occasional swollen neurons with pale, rarefied cytoplasm (black arrows); and corresponding semi-quantitative histopathological scoring (**F**) (n = 5). Data are presented as means ± s.d. Statistical significance was determined using log-rank Mantel-Cox test (**C**), two-way ANOVA with Tukey’s multiple comparisons test (**D**), or one-way ANOVA with Tukey’s multiple comparisons test (**E**, **G**). \**P* < 0.05; \*\**P* < 0.01; \*\*\**P* < 0.001; ns, not significant.The schematic in **A** was created with BioRender.com.

### Maternal LDL abundance limits ZIKV vertical transmission and congenital pathology

Given the potent anti-ZIKV activity of serum, we next asked whether this barrier varies under physiological conditions. Pregnancy provided a particularly relevant setting, as most maternal ZIKV infections are asymptomatic but fetal infection causes severe congenital abnormalities. We therefore investigated whether this intravascular antiviral barrier changes during pregnancy. Intact maternal LDL does not freely cross the placenta, whereas endogenous fetal apoB-containing lipoprotein production is limited and circulating LDL is rapidly utilized by growing fetal tissues^38^, resulting in substantially lower LDL abundance in the fetal circulation. To determine whether this physiological difference is accompanied by reduced antiviral activity, we analyzed paired third-trimester maternal and umbilical cord serum samples from 26 pregnancies. Cord blood LDL levels were only approximately one-eighth to one-tenth of maternal levels (**Fig. 7A**). Consistently, maternal serum exhibited strong antiviral activity, whereas cord blood serum showed significantly reduced antiviral capacity (**Fig. 7B**). These findings indicate that the LDL-dependent antiviral barrier is significantly attenuated in the fetal circulation and suggest that fetal protection depends primarily on maternal LDL intercepting ZIKV before it crosses the placenta.

**Fig. 7.**
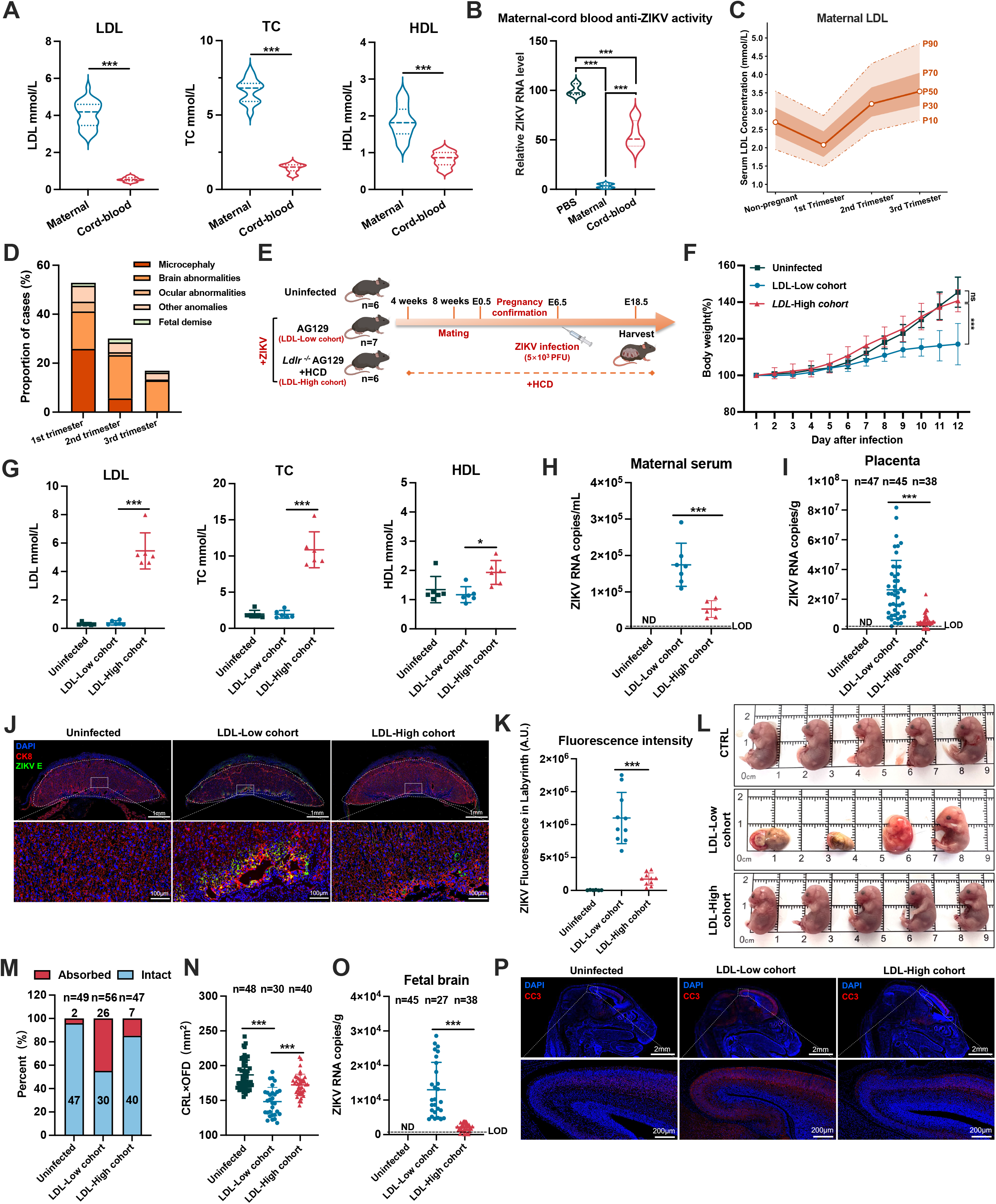
Maternal apoB-containing lipoproteins safeguard fetal development during ZIKV infection. **A**, LDL, TC and HDL concentrations in paired human maternal and umbilical cord-blood serum samples (n = 26 pairs). **B**, Huh7 cells were infected in the presence of 10% human maternal or cord-blood sera in **A** (n = 26). Relative intracellular ZIKV RNA levels at 36 hpi. were quantified. **C**, Serum LDL concentrations in non-pregnant women and across pregnancy trimesters reconstructed from published reference datasets. The orange line represents the central estimate, shaded areas indicate percentile-based reference ranges, and dashed lines mark the outer percentile limits. **D**, Proportion of abnormal fetal or neonatal outcomes associated with first-, second- or third-trimester maternal ZIKV infection across six published cohorts. Outcomes were classified as microcephaly, brain abnormalities, ocular abnormalities, other anomalies or fetal demise. **E**, Schematic diagram of the pregnant mouse infection model. AG129 dams were used as the LDL-low cohort, and *Ldlr*^−/-^ AG129 dams fed with a high-cholesterol diet were used as the LDL-high cohort. Pregnant dams were challenged subcutaneously with 5×10^3^ PFU ZIKV at embryonic day 6.5 (E6.5), and maternal, placental and fetal tissues were analysed at E18.5. Dam numbers were: uninfected, n = 6; ZIKV-infected LDL-low, n = 7; ZIKV-infected LDL-high, n = 6. **F**, Maternal body-weight changes from E6.5 to E18.5 (n = 6-7). **G**, Maternal serum LDL, TC and HDL concentrations at E18.5 (n = 6-7). **H**, Maternal serum ZIKV RNA levels at E18.5, quantified by Q-RT–PCR and expressed as RNA copies per mL (n = 6-7). **I**, Placental ZIKV RNA levels at E18.5, quantified by Q-RT–qPCR and expressed as RNA copies per gram of tissue (n = 38-45 placentas per group). **J**, **K**, Representative immunofluorescence staining of ZIKV E protein (red) and CK8 (green) in placental sections (**J**) and quantification of ZIKV E fluorescence intensity in the placental labyrinth zone (**K**; n = 10 placentas per group). **L**, **M**, Representative macroscopic images of intact maternal uteri at E18.5 (**L**) and quantification of fetal resorption (n = 47-56) (**M**). **N**, Fetal size at E18.5, quantified as crown–rump length (CRL) × occipitofrontal diameter (OFD) (n = 30 to 48). **O**, ZIKV RNA levels in fetal brains at E18.5, quantified by RT–qPCR and expressed as RNA copies per gram of tissue (n = 33-38). **P**, Representative immunofluorescence staining of cleaved caspase-3 in fetal brain sections. Data are presented as means ± s.d. Statistical significance was determined using mixed-effects analysis with Tukey’s multiple-comparison test (**F**), two-tailed unpaired Student’s *t*-test (**A, G**–**I**, **K**, **O**) or one-way ANOVA with Tukey’s multiple-comparison test (**B**, **N**). \**P* < 0.05; \*\*\**P* < 0.001; ns, not significant. The schematic in **E** was created with BioRender.com.

Maternal LDL abundance is not constant throughout pregnancy but reaches a physiological nadir during early gestation (**Fig. 7C**)^39–42^. To determine whether this period of reduced maternal LDL correlates with susceptibility to congenital ZIKV disease, we retrospectively analyzed six published pregnancy cohorts and case series from the 2015–2017 outbreak in the Americas, including studies from Brazil, the U.S. territories and the French territories of the Americas^43–48^. ZIKV-associated fetal or neonatal abnormalities, including microcephaly, brain abnormalities, ocular abnormalities and fetal demise, were reported following maternal infection during all three trimesters (**Fig. 7D** and **Fig. S7**). Notably, approximately 70% of all abnormal outcomes were associated with first-trimester maternal infection, which coincided with the physiological nadir of maternal LDL (**Fig. 7C, D** and **Fig. S7**). This temporal agreement in human suggests that reduced maternal LDL abundance during early pregnancy might weaken systemic viral interception and increase the possibility of ZIKV reaching the fetal compartment.

To establish the determinant role of maternal LDL in governing fetal infection risk, we therefore tested whether increasing maternal LDL could limit vertical transmission and congenital disease in vivo. Female *Ldlr*^−/-^ AG129 mice preconditioned with a high-cholesterol diet (HCD) were used as the maternal “LDL-High” cohort, with LDL levels approximating the upper range of LDL observed in human pregnancy (∼ 4.45 mmol/L), and female AG129 mice were raised as maternal “LDL-low” cohort. These female mice were subjected to timed mating and infected with ZIKV at embryonic day 6.5 (E6.5) (**Fig. 7E**). ZIKV infection severely restricted maternal weight gain in LDL-Low dams and this weight gain defect was attenuated in LDL-High dams **(Fig. 7F**). Serum analysis confirmed sustained elevations of LDL and total cholesterol in the LDL-High cohort (**Fig. 7G**). LDL-High dams displayed significantly attenuated viral burdens in maternal brain, spleen, and kidneys (**Fig. S8A–C**), which correlated with marked suppression of maternal viremia and placental viral dissemination (**Fig. 7H, I**). Histopathological examination revealed extensive disruption of the placental labyrinth zone in maternal LDL-Low mice, whereas structural integrity was better preserved in the LDL-High cohort (**Fig. S8D, E**). ZIKV envelope (ZIKV-E) antigen accumulation in the CK8-positive placental labyrinth zone of LDL-Low dams indicating placental infection; this infection was substantially reduced in maternal LDL-High cohort (**Fig. 7J, K**). Higher maternal LDL also reduced the extensive fetal resorption and growth restriction caused by ZIKV infection (**Fig. 7L, M**). Additionally, ZIKV-associated fetal growth restriction, reflected by reduced crown–rump length (CRL) and occipitofrontal diameter (OFD), was markedly alleviated in fetuses from the LDL-High cohort. (**Fig. 7N**). Fetal brain viral loads were significantly diminished (**Fig. 7O**) and cleaved caspase-3 staining in fetal brain sections was markedly reduced in the LDL-High cohort **(Fig. 7P)**. These data indicated that maternal LDL constituted innate immune barrier is a neglected determinant of ZIKV vertical transmission and congenital defects.

Together, results from maternal/cord blood analyses, retrospective human studies and congenital infection models coherently confirmed that maternal level of apoB-containing lipoproteins determines congenital ZIKV infection risk.

## DISCUSSION

In this study, we identified circulating apoB-containing lipoproteins as an unrecognized, dynamic intravascular innate immune barrier. By exposing surface-accessible phosphatidylserine (PS), these particles constitutively occupy PS-recognition receptors^7,10^, blocking ZIKV entry. Crucially, given the high abundance of LDL particles in human blood (∼10^15^ particles/mL)^49^, this receptor-competition mechanism provides a high-capacity extracellular buffer against apoptotic-mimicry viruses. By preventing receptor engagement and entry, this may prolong intravascular viral residence, allowing opsonization and clearance by the complement system^14,50^. Genetic or diet-induced elevation of apoB-containing lipoproteins enhanced serum antiviral activity and limited systemic ZIKV dissemination. In the gestational model, elevated maternal lipoproteins reduced maternal viremia, placental infection, fetal viral burden and fetal loss. These findings define apoB-containing lipoproteins as a dynamic intravascular innate immune barrier limiting viral entry, systemic dissemination and vertical transmission.

This LDL-dependent barrier may also intersect with other entry strategies^51^. Many viruses exploit lipoprotein pathways or receptors during entry, most notably hepatitis C virus through LDLR- and SR-BI-associated pathways^52,53^. Recent studies have also identified LDLR-family members as entry receptors for selected flaviviruses, including TBEV and yellow fever virus^3,54^. ApoB-containing lipoproteins may provide a bimodal defense: they may interfere with receptor pathways used by viruses that co-opt lipoprotein uptake machinery, while surface-exposed PS antagonizes apoptotic mimicry. In flavivirus-endemic regions, pre-existing dengue immunity can shape infection and disease risk through antibody-dependent enhancement (ADE)^55–57^. By facilitating viral entry via Fcγ receptors, ADE could enable the virion to bypass the PS-dependent LDL decoy shield. This mechanism may help explain the increased risk of ZIKV vertical transmission in DENV-immune pregnancies^58,59^, as the virus could evade both cross neutralization and the maternal lipid barrier.

Physiological variation in this barrier may also help explain the clinical dissociation between mild maternal infection and severe fetal disease. ZIKV infection in adults is usually asymptomatic or self-limited, whereas congenital abnormalities can follow mild or asymptomatic maternal infection. Fetal susceptibility may therefore depend not only on maternal disease severity but also on whether blood-borne virus escapes intravascular restriction and reaches the placenta. Maternal LDL reaches a physiological nadir (∼1.48 mmol/L) during early pregnancy^41,60^, overlapping with the gestational window most strongly associated with congenital abnormalities^46,61^. Our data support a model in which LDL decrease weakens the maternal-side intravascular barrier, permitting increased viremia and ZIKV seeding of the placenta. This vulnerability may be further amplified in the fetal circulation, where LDL levels are substantially lower than in maternal blood^62,63^. Serum from cord blood shows significantly weakened antiviral capability. After placenta penetration, LDL-poor blood may permit more efficient viral spread within susceptible fetal tissues. Therefore, high fetus ZIKV infection rate and following congenital defects may result from both transient decline in maternal LDL and the LDL-poor fetal compartment.

The vertical transmission model applied was established in AG129 mice which lack interferon responses. Although elevated maternal LDL reduced placental infection and fetal loss, ZIKV remained detectable in most fetuses. Clinically, approximately 70-80% of newborns from ZIKV infected mothers show no infection^46,64^, suggesting that complete pregnancy protection from ZIKV infection is dependent on the combination of apoB-containing lipoprotein barrier and innate immune response.

Overall, our work defines circulating apoB-containing lipoproteins as a blood-borne innate immune barrier. Following viral penetration through anatomic barrier, apoB-containing lipoproteins block the intravascular escape of ZIKV by continuously competing for PS-recognition pathways. This unrecognized function prevents prompt intravascular escape of viruses and reduces viral load during the time needed for complement opsonization, interferon response and adaptive immunity. Close monitoring of maternal LDL level of ZIKV contracted pregnant women is recommended to reduce vertical transmission possibility and to prevent following congenital infection.

## MATERIALS AND METHODS

### Cell lines

Human hepatocellular carcinoma cells (Huh7), human non-small cell lung cancer cells (A549), human embryonic kidney cells (293T), and African green monkey kidney cells (Vero E6) were kindly provided by B. Sun (Center for Excellence in Molecular Cell Science, Chinese Academy of Sciences). Human umbilical vein endothelial cells (HUVEC, CRL-1730) were obtained from the American Type Culture Collection (ATCC). All cells were maintained in Dulbecco’s Modified Eagle Medium (DMEM, Gibco/Invitrogen) supplemented with 10% heat-inactivated fetal bovine serum (FBS, Gibco), 1×non-essential amino acids, 100 U/mL penicillin, and 100 μg/mL streptomycin. Recombinant 293T cell lines stably expressing human TIM-1, AXL, DC-SIGN, or the TIM-1-WF/AA mutant were selected and maintained in complete DMEM containing 2 μg/mL puromycin. All cell lines used in this study were authenticated by short tandem repeat (STR) profiling and routinely verified to be free of mycoplasma contamination using PCR-based assays.

### Human serum samples

Serum samples were obtained from healthy adult donors at the Affiliated Hospital of Nantong University. Paired maternal and umbilical cord blood samples were collected at delivery. The study was approved by the institutional ethics committee of the Affiliated Hospital of Nantong University, and written informed consent was obtained from all participants. Serum was isolated after clotting, aliquoted, and stored at −80°C. Equal volumes of individual adult sera were pooled where indicated, and individual sera were used for LDL correlation analyses.

### Mice, dietary interventions, and ethical statement

Animal experiments complied with relevant ethical regulations and were approved by the Institutional Animal Care and Use Committee (IACUC) of the School of Basic Medical Sciences, Fudan University. Wild-type C57BL/6J, *Ldlr*^−/-^, and *Apoe*^−/-^ mice (fully backcrossed to a C57BL/6J background) were purchased from GemPharmatech (Nanjing, China). For the diet-induced obesity (DIO) model, C57BL/6J mice pre-fed a high-fat diet for 4 months was obtained from GemPharmatech. *Ifnar1*^−/-^*Ifngr1*^−/-^double-knockout (AG129) mice were sourced identically. Compound mutant AG129-*Ldlr*^−/-^ mice were custom-generated by GemPharmatech through crossbreeding. Mouse strain details and catalog numbers are listed in **Supplementary Table 1**. Animals were housed in a specific pathogen-free (SPF) facility (12-h light/dark cycle, 22 ± 2 °C, 50 ± 10% relative humidity) with *ad libitum* access to water and standard chow. AG129-*Ldlr*^−/-^ mice designated for dietary interventions were weaned at 4 weeks of age and transitioned to a customized high-cholesterol diet (HCD) (**Supplementary Table 2**).

### Viruses

Zika virus (ZIKV; GenBank KU321639.1) was derived from an infectious clone based on a strain isolated in Brazil. Dengue virus serotype 2 (DENV-2; strain 16681), West Nile virus (WNV; GenBank MZ595335), tick-borne encephalitis virus (TBEV; GenBank ON228408.1) and chikungunya virus (CHIKV; GenBank LN898093.1) were generated from infectious clones. Viral RNAs were *in vitro* transcribed (mMESSAGE mMACHINE T7 kit, Invitrogen) and electroporated into Vero E6 cells. Supernatants were harvested upon the appearance of cytopathic effects, clarified by centrifugation (700*g*, 5 min), filtered through a 0.22 μm filter, and stored at -80 °C. Hepatitis A virus (HAV, HM175/18f) and influenza A virus (IAV, H1N1 PR8) were prepared as previously described. Ebola virus-like particles were generated in Huh7-4PX cells using a reverse-genetics system provided by J. Zhong (Shanghai Institute of Immunity and Infection). SARS-CoV-2 (MT627325.1) was propagated in Vero E6 cells. Live SARS-CoV-2 experiments were conducted in a BSL-3 facility at the Naval Medical University.

### Serum-free ZIKV stock preparation

Serum-free ZIKV stocks were generated by propagating passage-1 ZIKV in Vero E6 cells. At 24 h post-infection, the culture medium was replaced with serum-free DMEM. Supernatants were collected at 48 h post-infection, clarified by centrifugation at 700g for 5 min, filtered through a 0.22 μm filter, aliquoted and stored at −80 °C. Viral titers were determined by plaque assay on Vero E6 cells. These serum-free stocks were used for serum and lipoprotein inhibition assays to avoid interference from FBS-derived components.

### Plasmid construction

Coding sequences of human *TIM-1* (NM_012206.3) and *DC-SIGN* (NM_021155.3) were cloned into the pWPI lentiviral vector using BamHI and MluI restriction sites. The phosphatidylserine (PS) binding-deficient mutant (TIM-1-WF/AA) was generated by introducing W112A and F113A mutations using the EasyGeno Homologous Recombination kit (Tiangen). For Fc-fusion proteins, an IL-2 signal peptide, and a human IgG1-Fc fragment were inserted into pWPI. The ectodomains of TIM-1 or TIM-1-WF/AA (amino acids P21–P290) were fused to the N-terminus of the IgG1-Fc sequence. All constructs were verified by Sanger sequencing.

### Viral titration

Infectious titers of ZIKV, WNV, TBEV, CHIKV, and DENV-2 were determined by plaque or focus-forming assays on Vero E6 cells. Monolayers in 24-well plates were inoculated with serial viral dilutions for 1 h at 37 °C, overlaid with 2% sodium carboxymethyl cellulose (CMC) in DMEM with 3% FBS, and incubated for 4 days. Cells were fixed with 4% paraformaldehyde (PFA). For ZIKV, WNV, TBEV, and CHIKV, cells were stained with 0.05% crystal violet, and macroscopic plaques were counted. For DENV-2, fixed cells were permeabilized, probed with a pan-flavivirus anti-E antibody (clone 4G2), and visualized using a fluorophore-conjugated secondary antibody. HAV titers were determined by an end-point dilution assay on Huh7.5.1-GA cells in 96-well plates; infected cells were quantified at 3 days post-infection by evaluating GFP nuclear translocation via fluorescence microscopy.

### Lipoprotein fractionation by density gradient ultracentrifugation

Serum lipoproteins were fractionated by discontinuous iodixanol density gradient ultracentrifugation. Serum (800 μl) was mixed with Opti Prep (60% w/v iodixanol; Serumwerk Bernburg) and underlaid in a 13.2-mL Ultra-Clear tube (Beckman Coulter). A discontinuous gradient was formed by sequentially overlaying 2 mL each of 80%, 60%, 40%, and 20% Opti Prep solutions (v/v in PBS), topped with PBS. Samples were centrifuged at 154,000*g* for 16 h at 4 °C in an SW 41 Ti rotor (Optima L-100 XP, Beckman Coulter). Acceleration was set to “slow,” and the brake was disabled. Eleven fractions were sequentially collected from the top. Fraction densities were verified via refractive index measurements using an Abbe refractometer.

### ApoB-containing lipoprotein depletion from human serum

ApoB-containing lipoproteins were depleted from human serum by antibody-mediated immunocapture. Protein G magnetic beads (400 μl; Thermo Fisher Scientific) were washed three times with PBS and incubated with 50 μg anti-ApoB antibody or species-matched control IgG for 8 h at 4 °C with gentle rotation. Antibody-coupled beads were washed three times with PBS to remove unbound antibody and then incubated with 20 μl human serum for 12 h at 4 °C with gentle rotation. The control-depletion condition used the same bead volume, antibody amount, incubation time and temperature. Beads were separated using a magnetic rack, and the unbound serum fraction was collected as ApoB-depleted or control-depleted serum. Depletion efficiency was determined by immunoblotting for ApoB. For ZIKV inhibition assays, ApoB-depleted and control-depleted sera were diluted into infection medium at a final concentration of 10% (v/v).

### Lipoprotein preparation and biochemical modification

Commercially purified human native LDL (20613ES), VLDL (20617ES), and oxidized LDL (oxLDL; 20605ES) were purchased from Yeasen Biotechnology (Shanghai, China). To enzymatically deplete surface-exposed phosphatidylserine (PS), 50 μg of VLDL, native LDL or oxidized LDL was incubated with 100 U of phospholipase D (PLD; MedChemExpress, HY-P2812) at 37 °C for 1 h. The enzymatic reaction was subsequently terminated by the addition of EDTA (final concentration 10 mM). Following modification, accessible surface PS levels on all lipoprotein variants were quantitatively determined utilizing a commercial PS ELISA kit (ELK Biotechnology, ELK8149) according to the manufacturer’s protocols.

### Expression and purification of recombinant Fc-fusion proteins

Lentiviral-transduced 293T cells stably expressing Fc-fusion constructs were expanded in 15-cm dishes. At 90% confluence, cells were washed with PBS and transitioned to serum-free 293 Expression Medium (Gibco). Supernatants were harvested 72 h later, clarified (700*g* for 5 min; 2000*g* for 20 min), and concentrated using 10-kDa MWCO ultrafiltration units (Amicon Ultra-15, Millipore). Proteins were purified using Protein G 4FF agarose beads (AOGOMA), eluted with 0.1 M glycine (pH 2.8), and immediately neutralized with 1 M Tris-HCl (pH 9.0). Purified proteins were buffer-exchanged into PBS, quantified via BCA assay (Beyotime), evaluated by SDS–PAGE, and stored at -80 °C.

### Preparation of artificial liposomes

Lipid mixtures of pure phosphatidylcholine (PC) or a PS/PC combination (3:7 molar ratio) in chloroform were evaporated under reduced pressure at 30 °C. Lipid films were desiccated under high vacuum for 1 h, hydrated in PBS (pH 7.4) to 10 mM total lipid, and dispersed by vortexing. Suspensions were sonicated on ice for 5 min to form small unilamellar vesicles. Large aggregates were pelleted at 10,000*g* for 10 min, and the clarified liposome supernatants were stored at -80 °C.

### CRISPR/Cas9-mediated gene knockout and siRNA silencing

To establish *TIM-1* knockout cells, A549 and Huh7 cells were transduced with Lenti-CRISPR v2 particles encoding Cas9 and a *TIM-1*-specific sgRNA (5’-CACCGTGGCAGGGTAGTGTGACAGA-3’). Cells were selected with 1 μg/mL puromycin for 7 days and single-cell sorted into 96-well plates. Knockout efficiency of clonal populations was validated by immunoblotting. For transient silencing, Huh7 cells were transfected with 20 pmol of siRNAs targeting *SR-BI*, *SDC-1*, *LDLR*, or *TIM-1* using Lipofectamine RNAiMAX (Invitrogen). Knockdown efficiency was quantified by RT-qPCR 24 h post-transfection. The specific siRNA sequences (Forward; Reverse) utilized were as follows:

SR-BI-F: UCAUGAUGAGCUUCAGGGUCAUGGG;

SR-BI-R: CCCAUGACCCUGAAGCUCAUCAUGA;

SDC-1-F: AUUAGUAGCCACAAUUUGCGGCAGG;

SDC-1-R: CCUGCCGCAAAUUGUGGCUACUAAU;

LDLR-F: UCGUUGAUGAUAUCUGUCCAAAAUA;

LDLR-R: UAUUUUGGACAGAUAUCAUCAACGA;

TIM-1-F: UGUAGUGGCAGGGUAGUGUGACAGA;

TIM-1-R: UCUGUCACACUACCCUGCCACUACA;

AXL-F: GGCUCUCCAAGAAGAUCUACA;

AXL-R: UAGAUCUUCUUGGAGAGCCCG;

Non-targeting control-F: ACGUGACACGUUCGGAGAATT;

Non-targeting control-R: UUCUCCGAACGUGUCACGUTT.

### Viral attachment assays

Confluent Huh7 monolayers were pre-chilled at 4 °C for 45 min, followed by co-incubation with ZIKV (MOI = 10) and 150 μg/mL of BSA, VLDL, or LDL in cold, serum-free DMEM for 1 h at 4 °C. Monolayers were washed three times with cold PBS. Surface-bound virions were quantified by extracting total RNA using TRIzol Universal Reagent (Tiangen) and analyzing ZIKV genome copies via RT-qPCR.

### In vitro pull-down and competitive binding assays

Purified human IgG1-Fc, TIM-1-Fc, or TIM-1-WF/AA-Fc (10 μg) were immobilized on 10 μl of Protein G magnetic beads (Thermo Fisher Scientific) in PBS for 2 h at 4 °C. Beads were washed with PBST (PBS with 0.02% Tween-20) and incubated with either 5×10^6^ PFU of ZIKV or 20 μg of purified VLDL/LDL in a 500 μL reaction volume for 12 h at 4 °C. For competitive binding, TIM-1-Fc-functionalized beads were co-incubated with 5×10□ PFU of ZIKV and 20 μg of VLDL, LDL, or BSA. Beads were washed six times with PBST. Captured complexes were eluted in 1× Laemmli buffer at 95 °C for 10 min, resolved by SDS-PAGE, and analyzed by immunoblotting.

### Annexin V masking assay

Purified LDL (50 μg/mL) was pre-incubated with 1–10 μg/mL recombinant Annexin V (Abcam, ab89493) in binding buffer (10 mM HEPES, 140 mM NaCl, 2.5 mM CaCl□, pH 7.4) for 30 min at room temperature prior to DiI-LDL uptake or ZIKV infection assays. Annexin V-free buffer served as the vehicle control.

### ZIKV subgenomic replicon assay

In vitro-transcribed RNAs of the ZIKV subgenomic replicon (ZIKV-SGR, expressing a Renilla luciferase reporter) or its polymerase-deficient ΔGDD mutant were electroporated (10 μg RNA per 4 × 10□ cells) into Huh7 cells (Bio-Rad Gene Pulser; 270 V, 950 μF). At 6 h post-transfection, cells were treated with 150 μg/mL BSA, VLDL, or LDL. Viral replication and translation kinetics were assessed at indicated time points by quantifying luciferase activity (Renilla-Glo, Promega) and RT–qPCR.

### DiI-LDL uptake assay

Cells (5×10^4^ cells/well) were incubated with 5 μg/mL DiI-labeled LDL (Yeasen, 20614ES76) for 2 h at 37 °C. After washing with PBS, cells were fixed with 4% PFA and counterstained with Hoechst 33342 (Sangon Biotech). Fluorescence images were acquired using an Olympus IX53 microscope. Mean fluorescence intensity was quantified across six random fields per sample using ImageJ.

### ZIKV infection in adult mouse models

Eight-week-old male mice were allocated to four groups: mock-infected AG129, ZIKV-infected AG129, ZIKV-infected AG129-*Ldlr*^−/-^, and HCD-fed ZIKV-infected AG129-*Ldlr*^−/-^. Mice were inoculated subcutaneously (s.c.) with 5×10³ PFU of ZIKV. Experiments were performed in two independent cohorts. One cohort was monitored daily for survival up to 14 days post-infection (dpi). A second cohort was monitored for body weight changes and euthanized at 10 dpi. Plasma was isolated for lipid profiling and viremia quantification. Target organs (brain, liver, spleen, kidney) were harvested for viral RNA quantification. Brains were fixed for H&E staining.

### Timed mating and gestational ZIKV infection

Female mice (8-10 weeks old) in estrus were co-housed overnight with single stud males (2:1 ratio). The presence of a vaginal plug the following morning was designated embryonic day 0.5 (E0.5). At E6.5, pregnant dams were inoculated s.c. with 5×10³ PFU of ZIKV. Dams were euthanized at E18.5, and intact uteri were photographed *in situ*. Maternal plasma, brain, spleen, and kidneys were collected for lipid analysis, viral RNA quantification, and histology.

### Fetal biometry and pathology

Placentas and fetuses were dissected from the uterine horns. Fetal resorption was defined macroscopically by hemorrhagic remnants or arrested conceptuses measuring <1 cm. The fetal resorption rate was calculated as the ratio of resorbed conceptuses to the total number of implantation sites per litter. Crown-rump length (CRL) and occipitofrontal diameter (OFD) of viable fetuses were measured using a tissue measurement board. Fetal heads and placentas were homogenized for ZIKV RNA quantification. Placentas were processed for H&E staining and dual immunofluorescence (co-staining for ZIKV E and CK8 or Vimentin). Fetuses were stained for Cleaved-Caspase 3.

### RNA extraction and quantitative real-time PCR (RT-qPCR)

Total RNA was extracted from cells or supernatants using TRIzol Universal Reagent (Tiangen) and quantified (NanoDrop 2000). RNA was reverse transcribed using the ReverTra Ace qPCR RT Kit (Toyobo). RT-qPCR was performed on an ABI Quant Studio 6 Flex system with SYBR Green Master Mix (Toyobo). Absolute quantification of ZIKV was determined using standard curves generated from *in vitro*-transcribed viral RNA (10^4^ to 10^10^ genome equivalents). Intracellular viral loads were normalized as genome equivalents (GE) per μg of total RNA; extracellular titers were expressed as GE/mL. Relative host gene expression was calculated using the 2^−ΔΔCt^ method, normalized to *GAPDH*. Primer sequences are listed in **Supplementary Table 3**.

### Plasma lipid profiling

Plasma concentrations of total cholesterol (TC), high-density lipoprotein cholesterol (HDL), and low-density lipoprotein cholesterol (LDL) were quantified using commercial enzymatic colorimetric assay kits (Mindray, Shenzhen, China). TC levels were measured utilizing the standard cholesterol oxidase method. HDL and LDL fractions were determined via homogeneous direct clearance methods. Automated lipid biochemical assays were performed on a Mindray BS-2800M clinical chemistry analyzer.

### Analysis of pregnancy-associated LDL dynamics

Published reference datasets reporting serum LDL-C concentrations in non-pregnant and pregnant women were curated from four studies. Data were extracted by gestational stage and harmonized into four groups: non-pregnant, first trimester, second trimester and third trimester. When values were reported in mg dl□¹, they were converted to mmol l□¹ using a conversion factor of 38.67. For each stage, the median or closest available central estimate was used to define the main trajectory and reported percentile, or interquartile limits were used to define shaded reference intervals. Adult non-pregnant LDL-C thresholds were derived from the Chinese Guidelines for Lipid Management 2023 and the 2018 AHA/ACC cholesterol guideline.

### Synthesis of published pregnancy cohorts with trimester-resolved maternal ZIKV infection

Published studies reporting fetal or neonatal abnormalities after maternal ZIKV infection were identified through PubMed/MEDLINE searches and reference screening. Studies were included if they reported maternal ZIKV infection or exposure, abnormal fetal or neonatal outcomes, and the trimester of maternal infection. Cohort studies, case-control studies, case series, and surveillance reports were included. Animal and in vitro studies, reviews, guidelines, studies lacking trimester-specific infection timing or abnormal fetal or neonatal outcomes, and overlapping cohorts were excluded. For overlapping populations, the larger or more informative study was retained.

The timing of maternal infection was assigned according to gestational age at symptom onset, rash onset, molecular diagnosis, or reported exposure, as defined in each study. When more than one measure was available, symptom or rash onset was used preferentially. Cases with unknown timing, periconceptional exposure, or exposure spanning more than one trimester were excluded unless trimester-specific assignment was reported. Outcomes included microcephaly, brain or ocular abnormalities, other congenital anomalies, congenital Zika syndrome or Zika-associated birth defects, fetal loss, miscarriage, stillbirth, and termination because of fetal abnormalities.

For each study, the proportion of affected fetuses or infants was calculated for maternal infection occurring in the first, second, or third trimester among cases with known infection timing. Trimester-specific proportions were summarized using random-effects models, and between-study heterogeneity was assessed using I².

### Immunofluorescence microscopy

ZIKV-infected monolayers were fixed with 4% PFA for 15 min, permeabilized with 0.5% Triton X-100 for 10 min and blocked with 5% FBS. Cells were probed with a mouse anti-ZIKV envelope antibody (clone 4G2, Thermo) for 1 h. After washing, samples were incubated with an Alexa Fluor 488-conjugated anti-mouse secondary antibody (Abclonal) for 45 min. Nuclei were counterstained with DAPI (Beyotime). Digital images from six fields per sample were captured using an Olympus IX53 microscope; the percentage of ZIKV-positive cells was quantified using Image-Pro Plus.

### Immunoblotting analysis

Protein samples in 1× Laemmli buffer were denatured at 98 °C for 5 min, resolved by SDS-PAGE, and transferred to PVDF membranes. After blocking with 5% non-fat milk in TBST (0.1% Tween-20) for 1 h, membranes were incubated overnight at 4 °C with primary antibodies targeting ZIKV E, Core, and NS5 (laboratory-generated polyclonal antibodies), APOB (Abclonal A4184), APOE (Abclonal 16344), APOA1 (Abclonal A1129), GAPDH (Abclonal AC033), the human Fc domain (Abclonal A21291), TIM-1 (R&D Systems AF1727), or AXL (Abclonal A17874). Membranes were washed and probed with HRP-conjugated secondary antibodies. Signals were visualized using ECL reagent (Bio-Rad) and captured on an E-Bolt imaging system.

### Histopathology and immunofluorescence microscopy

Excised placentas and maternal brains were fixed in universal tissue fixative for at least 24 h, dehydrated, and embedded in paraffin. Paraffin blocks were sectioned at 4 μm thickness. For histopathology, sections were deparaffinized, rehydrated, and stained with H&E. Maternal brain and placental sections were blindly scored by an independent commercial laboratory (Servicebio, Wuhan, China). ZIKV-induced cytopathic effects—specifically, neuronal shrinkage, hyperchromasia, and loss of nucleocytoplasmic demarcations in the brain, and trophoblast barrier disruption in the placenta—were semi-quantitatively graded from 0 to 4 based on the affected tissue area: 0, no lesions; 1, <25%; 2, 25–50%; 3, 50–75%; and 4, >75%.

For immunofluorescence, sections were subjected to heat-induced antigen retrieval using citrate buffer (pH 6.0) or Tris-EDTA buffer and blocked with 3% BSA. Sections were incubated overnight at 4 °C with primary antibodies targeting the ZIKV envelope protein (Sino Biological, 40543-MM09), Cytokeratin 8 (Servicebio, GB12233), Vimentin (Servicebio, GB15192), or Cleaved-Caspase 3 (Servicebio, GB11532). Sections were probed with corresponding Alexa Fluor 488- or 594-conjugated secondary antibodies (Servicebio) and counterstained with DAPI. Slide digitization was performed utilizing an LG-S80 slide scanner (Servicebio).

### Study design, randomization, and blinding

For *in vivo* studies, age- and sex-matched mice were randomly assigned to experimental or control dietary/infection groups. Adult mechanistic models utilized male mice to exclude the confounding variables of estrous cycle-induced lipid fluctuations, whereas gestational models exclusively utilized female mice. Investigators responsible for conducting *in vivo* viability assessments, measuring fetal biometrics (CRL and OFD), and performing histopathological scoring were completely blinded to the maternal genotype and dietary group allocation until the final data lock.

### Statistical analysis

All statistical analyses were performed using GraphPad Prism software (version 10.4.0) or R (version 4.5.2). Statistical analyses were conducted as described in the figure legends. Comparisons between two independent groups were analyzed using two-tailed unpaired Student’s t-tests. For experiments involving three or more groups, data were analyzed using one-way analysis of variance (ANOVA) followed by Dunnett’s post-test when comparing experimental groups to a single control, or Tukey’s post-test when comparing all groups against each other. Datasets with two independent variables were evaluated using two-way ANOVA with Šidák’s post-test. Repeated measures across time points were assessed using a mixed-effects analysis followed by Tukey’s post-test. Survival curves were compared using the log-rank Mantel-Cox test. Relationships between continuous variables were determined using simple linear regression with Pearson correlation. Statistical significance is defined as \**P* < 0.05, \*\**P* < 0.01, and \*\*\**P* < 0.001. Non-significant differences are denoted as ns.

## Data availability

For original data, please contact

## Acknowledgments

We thank Dr. Jinghua Yan (Institute of Microbiology, Chinese Academy of Sciences) for providing Z23 ZIKV neutralizing antibodies and Dr. Ping Zhao (Department of Microbiology, Faculty of Naval Medicine, Naval Medical University, Shanghai, China) for providing SARS-COV-2. This work was supported by China-Germany collaborative project from National Natural Science Foundation of China (NSFC) and Deutsche Forschungsgemeinschaft (DFG) (NSFC-DFG 81761138046 to Gang Long and Ralf Bartenschlager), Shanghai Municipal Science and Technology Major Project (ZD2021CY001) and NSFC general projects (NSFC 82372253, NSFC 82572568 to Gang Long).

## Author contributions

J.C., D.L. and G.L. conceived the study and designed the experimental strategy. J.C., D.L., W.C., Y.C., Y.Z., N.L., N.D., L.G., X.J., J.Q., G.Z., M.X. and Y.T. performed the investigation. J.C., D.L., W.C., Y.C. and G.L. developed the methodology, performed validation and analysed the data. J.Z. and Y.T. provided resources. R.B. contributed to fund raising and manuscript review. J.C. and G.L. wrote the original draft. J.C., W.C., D.L., R.B. and G.L. reviewed and edited the manuscript. G.L. curated the data, supervised the project and acquired funding. All authors discussed the results and approved the final manuscript.

## Competing interests

The authors declare no competing interests.

## Notes

### Competing Interest Statement

The authors have declared no competing interest.

