## Supplemental tables and figures for "Circulating apoB-containing lipoproteins constituted blood-borne innate immune barrier limits ZIKV dissemination and congenital infection risk": Supplementary Dataset.pdf

### Supplement tables and figures

**Supplemental table 1. Primers for quantitative real-time PCR**

| Target | Sense sequence (5'–3') | Antisense sequence (5'–3') |
|---|---|---|
| ZIKV | GTTTGGCTGGCCTATCAGGT | GTGTCTGGTCCACACCTCTG |
| WNV | AGGCCGGGTAGAGATTGACT | ACTTTCCGCTCTCTGTGGTG |
| TBEV | AAGCCCTGTAGGATCCCAGT | TGAAAGCACCTCCAAGGACC |
| DENV-2 | TCAAGGCTGAGAGGGTTATAGA | TTAGAGTGGGTCACTGGCAT |
| CHIKV | AAGCTCCGCGTCCTTTACCAA | ATTTAGCGTCCTTAACTGTGACG |
| HAV | GGTAGGCTACGGGTGAAAC | AACAACTCACCAATATCCGC |
| HSV-1 | AACGCGACGCACATCAAG | CTGGTACGCGATCAGAAAGC |
| *SR-BI* | GGTCCAGAACATCAGCAGGATC | GCCACATTTGCCCAGAAGTTCC |
| *SDC-1* | TCCTGGACAGGAAAGAGGTGCT | TGTTTCGGCTCCTCCAAGGAGT |
| *LDLR* | GAATCTACTGGTCTGACCTGTCC | GGTCCAGTAGATGTTGCTGTGG |
| *TIM-1* | CTTCACCTCAGCCAGCAGAAAC | GCCATCTGAAGACTCTGTCACG |
| *GAPDH* | GAAGGTGAAGGTCGGAGTC | GAAGATGGTGATGGGATTTC |

**Supplemental table 2. Mouse strains, genetic backgrounds, and commercial sources.**

| Mouse strains | Catalog numbers |
|---|---|
| *C57BL/6JGpt* | N000013 |
| *C57BL/6JGpt DIO* | T002040 |
| *C57BL/6JGpt* Ldlr$^{-/-}$ | T001464 |
| *C57BL/6JGpt* Apoe$^{-/-}$ | T001458 |
| *C57BL/6JGpt Ifnar1*$^{-/-}$-*Ifngr1*$^{-/-}$ | T055933 |

**Supplemental table 3. Composition of the matched purified experimental diets.**

| Product | Normal diet | | High cholesterol diet | |
|---|---|---|---|---|
| | gm% | kcal% | gm% | kcal% |
| Protein | 19.2% | 20% | 26% | 20% |
| Carbohydrate | 66.7% | 70% | 25% | 20% |
| Fat | 4.3% | 10% | 34% | 60% |
| Total | | 100% | | 100% |
| kcal/gm | 3.82 | | 5.15 | |
| **Ingredient** | **gm** | **kcal** | **gm** | **kcal** |
| Casein | 200 | 800 | 200 | 800 |
| L-Cystine | 3 | 12 | 3 | 12 |
| Cornstarch | 506.20 | 2024.8 | 0 | 0 |
| Maltodextrin | 125 | 500 | 125 | 500 |
| Sucrose | 72.8 | 291.2 | 72.8 | 291.2 |
| Cellulose, BW200 | 50 | 0 | 50 | 0 |
| Soybean Oil | 25 | 225 | 25 | 225 |
| Lard | 20 | 180 | 245 | 2205 |
| Mineral Mix S10026B | 50 | 0 | 50 | 0 |
| Vitamin Mix V10001C | 1 | 4 | 1 | 4 |
| Choline Bitartrate | 2 | 0 | 2 | 0 |
| Cholesterol | 0 | 0 | 9.8 | 0 |
| FD&C Yellow Dye#5 | 0.05 | 0 | 0 | 0 |
| FD&C Blue Dye#1 | 0 | 0 | 0.05 | 0 |
| Total | 1055.05 | 4037.0 | 783.64557 | 4037.2 |

**Supplemental table 4. PCR Primers for validation of triple-deficient Ldlr-/-AG129 mice**

| Target | Sense sequence (5'–3') | Antisense sequence (5'–3') |
|---|---|---|
| AF/R1 | CGTGGAGACTGAGCTGAGATGAAG | CACTGTGTGTGCTGGAAATGACAG |
| AF/R2 | CTCCCAGGATGACTTCCGAT | CGCAGTGCTCCTCATCTGAC |
| GF/R1 | CACAATCTCAGGGGTGCAATCTATC | ACGTTAGAAAGACAGACATGCGC |
| GF/R2 | TCATGTTGTGGTGATCCTAGCC | CAGGGAGGTCTCAGACACTTAAAC |
| LF/R | CTCCCAGGATGACTTCCGAT | CGCAGTGCTCCTCATCTGAC |

### Supplemental Figures and Legends

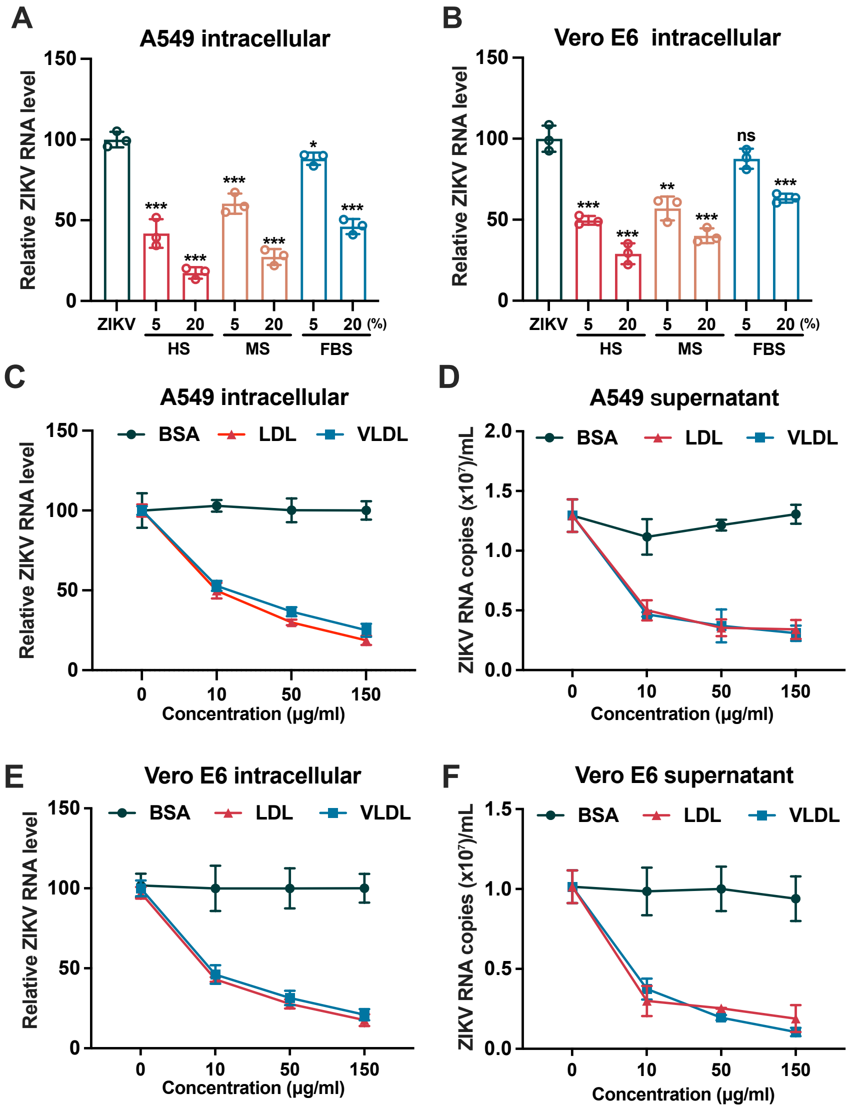


**Supplemental figure 1. Serum and apoB-containing lipoproteins restrict ZIKV infection across cell types.**

**A, B,** Relative intracellular ZIKV RNA levels normalized to GAPDH, measured by Q-RT–qPCR in A549 (**A**) and Vero E6 (**B**) cells at 24 hours post-infection (hpi). Cells were infected with ZIKV (MOI = 0.1) in media supplemented with 20% (v/v) fetal bovine serum (FBS), pooled mouse serum (MS), or pooled human serum (HS). **C–F,** Relative intracellular ZIKV RNA levels normalized to GAPDH (**C, E**), and extracellular viral RNA (**D, F**) quantified by Q-RT-qPCR in A549 (**C, D**) and Vero E6 (**E, F**) cells at 36 hpi. Cells were infected (MOI = 0.1) in the presence of indicated concentrations of bovine serum albumin (BSA), VLDL or LDL. Data are presented as means ± s.d. from three independent experiments. Statistical analysis was performed using one-way ANOVA with Dunnett's post-test (**A–F**). *$P < 0.05$; **$P < 0.01$; ***$P < 0.001$; ns, not significant.

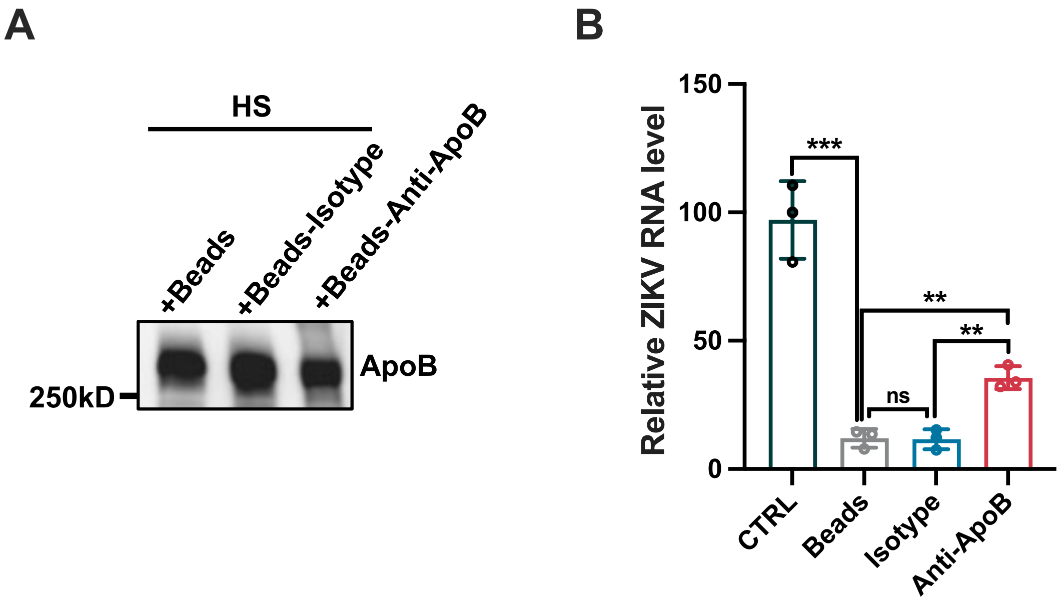


**Supplemental figure 2. Partial depletion of ApoB-containing lipoproteins reduces ZIKV restriction capability of human serum.**

**A,** Immunoblot analysis of APOB and APOE in human serum after control IgG or anti-ApoB antibody-mediated depletion. **B,** Relative intracellular ZIKV RNA levels in Huh7 cells infected in the presence of control-depleted or ApoB-depleted human serum. Data are presented as means ± s.d. from three independent experiments. Statistical analysis was performed using one-way ANOVA with Tukey's multiple-comparison test (**B**). **$P < 0.01$; ns, not significant.

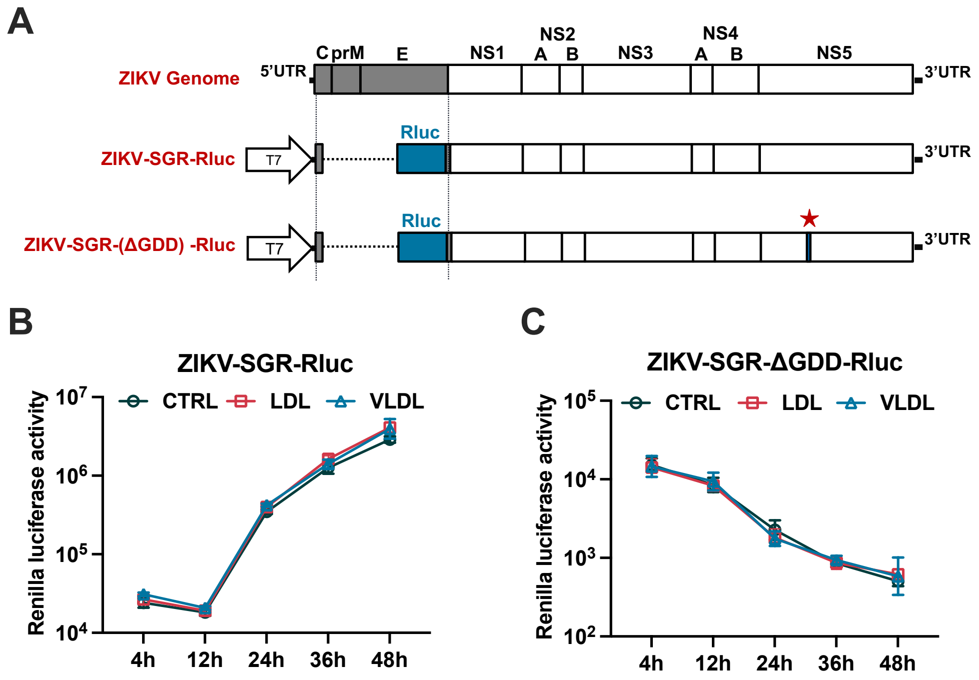


**Supplemental figure 3. ZIKV intracellular replication and translation are unaffected by apoB-containing lipoproteins.**

**A,** Schematic representation of the wild-type ZIKV subgenomic replicon (ZIKV-SGR) and the polymerase-defect mutant (ZIKV-SGR ΔGDD) constructs. **B, C,** Luciferase activity was measured in Huh7 cells transfected with ZIKV-SGR (**B**) or ZIKV-SGR ΔGDD (**C**). At 6 h post-transfection, the culture medium was replaced with fresh medium containing 150 μg/mL VLDL or LDL. Assays were performed at indicated time points post-transfection. Data are presented as means ± s.d. from three independent experiments. Statistical significance was determined using mixed-effects analysis with Tukey's multiple-comparison test (**B**, **C**).

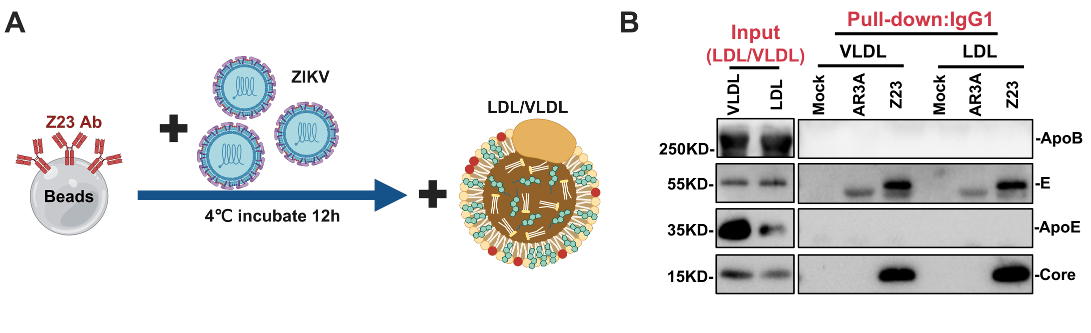


**Supplemental figure 4. ApoB-containing lipoproteins do not directly bind ZIKV particles.**

**A**, Schematic of the *in vitro* viral immunoprecipitation assay. **B**, Immunoblot analysis of complexes captured by Protein G magnetic beads coated with 10 µg of an HCV neutralizing antibody (AR3A, negative control) or a ZIKV neutralizing antibody (Z23), following co-incubation with 5 × $10^6$ PFU of ZIKV and 20 µg of purified VLDL or LDL.

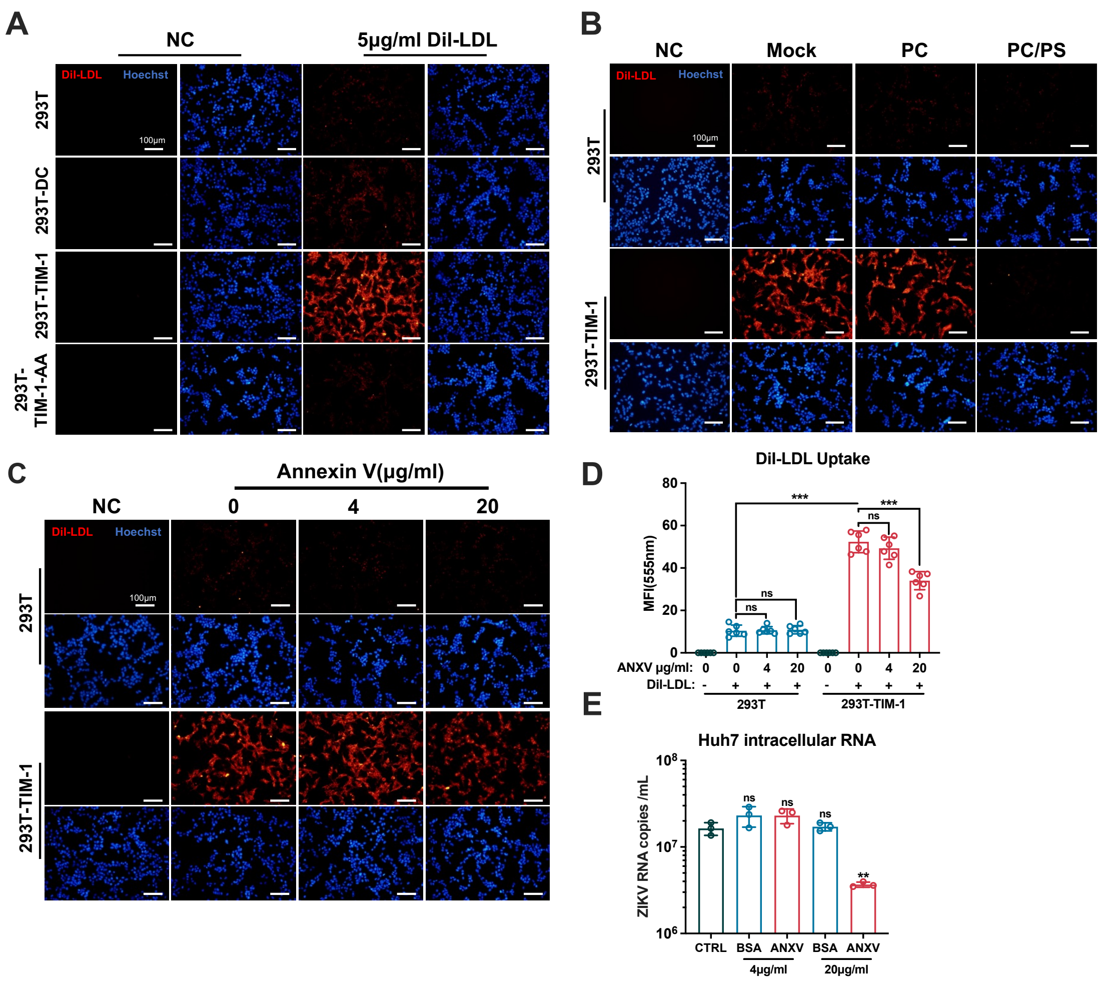


**Supplemental figure 5. ApoB-containing lipoproteins engage PS-recognition receptors through surface-exposed PS.**

**A–C,** Representative Hoechst-counterstained fluorescence images of DiI-LDL internalization (red). DiI-LDL uptake was assessed following a 2-h incubation with 5 µg/mL DiI-LDL in wild-type 293T cells and cells expressing DC-SIGN, wild-type TIM-1, or the PS-binding-deficient TIM-1-WF/AA mutant (**A**). DiI-LDL uptake in 293T-TIM-1 cells was further evaluated in the presence of 20 µM PC or PC/PS liposomes (**B**), or the indicated concentrations of recombinant Annexin V (**C**). **D,** Quantification of mean DiI-LDL fluorescence intensity from individual cells (n = 6) under the Annexin V blocking conditions was shown in **C**. **E,** Intracellular ZIKV RNA levels in Huh7 cells at

24 h post-infection (hpi). ZIKV (MOI = 0.1) was pre-incubated with the indicated concentrations of bovine serum albumin (BSA, control) or Annexin V for 2 h prior to inoculation. Data are presented as means ± s.d. from three independent experiments. Statistical significance was determined using two-way ANOVA with Šídák's multiple comparisons test (**D**) or one-way ANOVA with Dunnett's multiple comparisons test (**E**). $^{**}P < 0.01$; $^{***}P < 0.001$; ns, not significant.

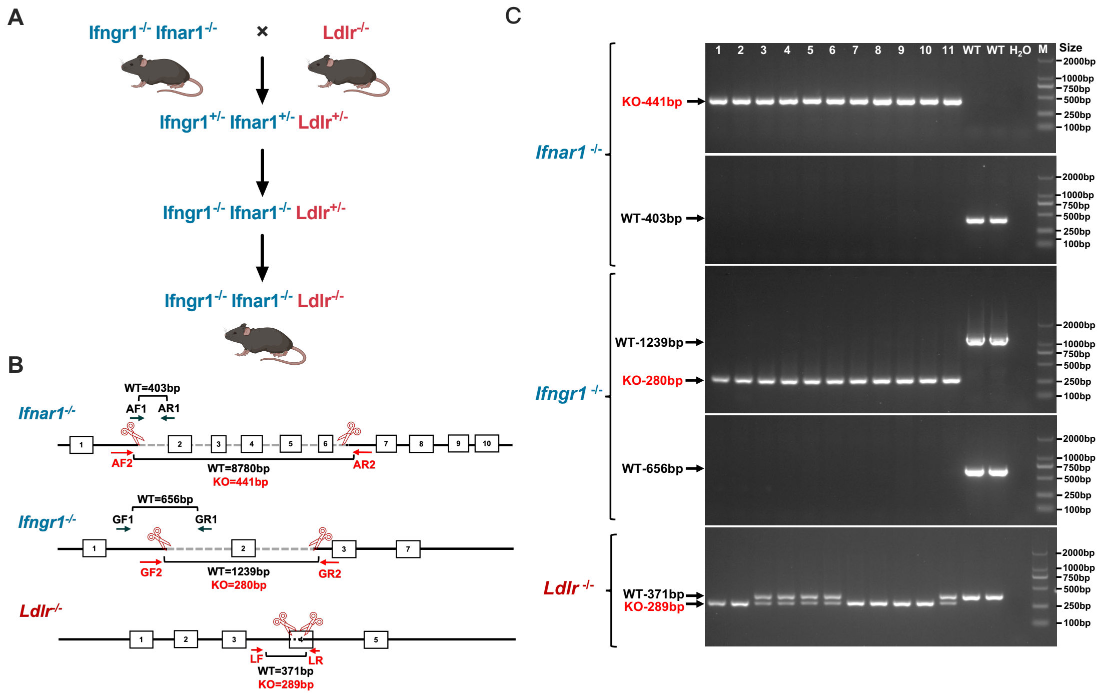


**Supplemental figure 6. Generation and validation of triple-deficient AG129-*Ldlr*$^{-/-}$mice.**

**A**, Schematic diagram of the breeding strategy utilized to generate AG129-*Ldlr*$^{-/-}$ (triple-knockout) mice on a C57BL/6J background. **B**, Strategy for genomic deletion validation, including the target DNA fragment and the specific binding sites of the genotyping primers (Supplement table 4). **C**, PCR genotyping results for the *Ifnar1*, *Ifngr1*, and *Ldlr* loci, confirming the successful construction of the triple-deficient mouse strain.

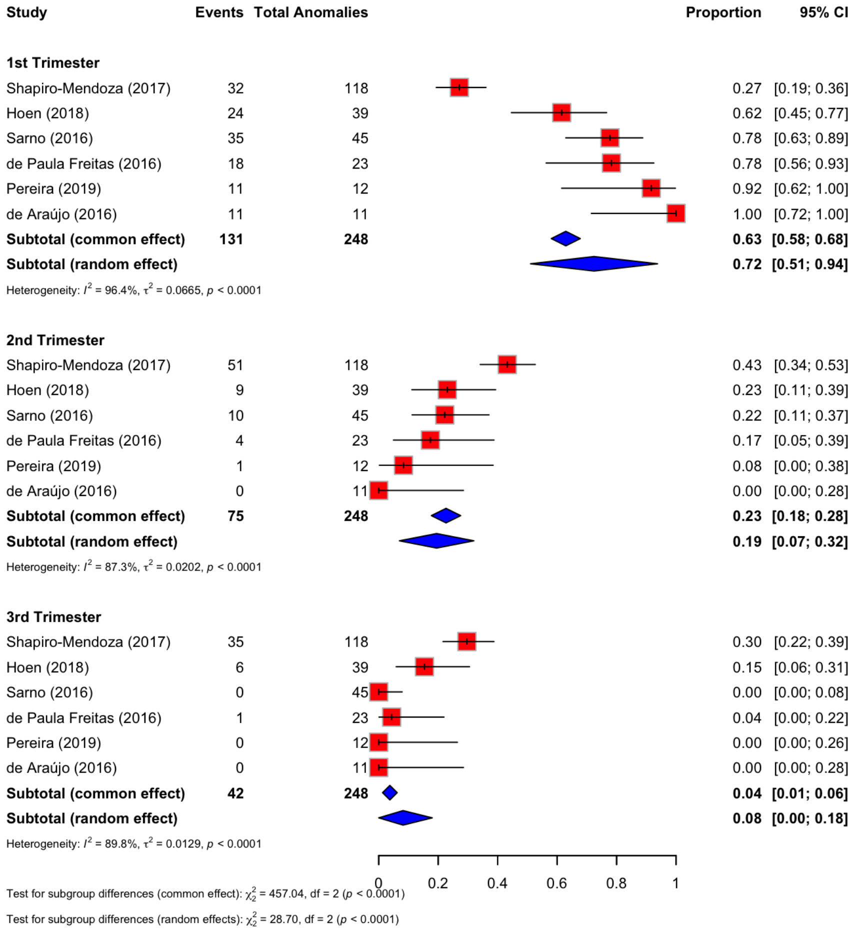


**Supplemental figure 7. Analysis of gestational timing of maternal ZIKV infection among abnormal pregnancy outcomes.**

Forest plots showing the proportion of affected fetuses or infants whose mothers had ZIKV infection, symptoms or reported exposure during the first, second or third trimester across six published cohorts or case series. Only fetuses or infants with congenital anomalies, Zika-associated birth defects or fetal demise and known maternal infection timing were included. Cases with unknown timing, periconceptional exposure or exposure spanning multiple trimesters were excluded unless trimester-specific allocation was explicitly available. Pooled trimester-specific proportions were estimated using random-effects models as a descriptive synthesis. Heterogeneity was assessed using $I^2$.

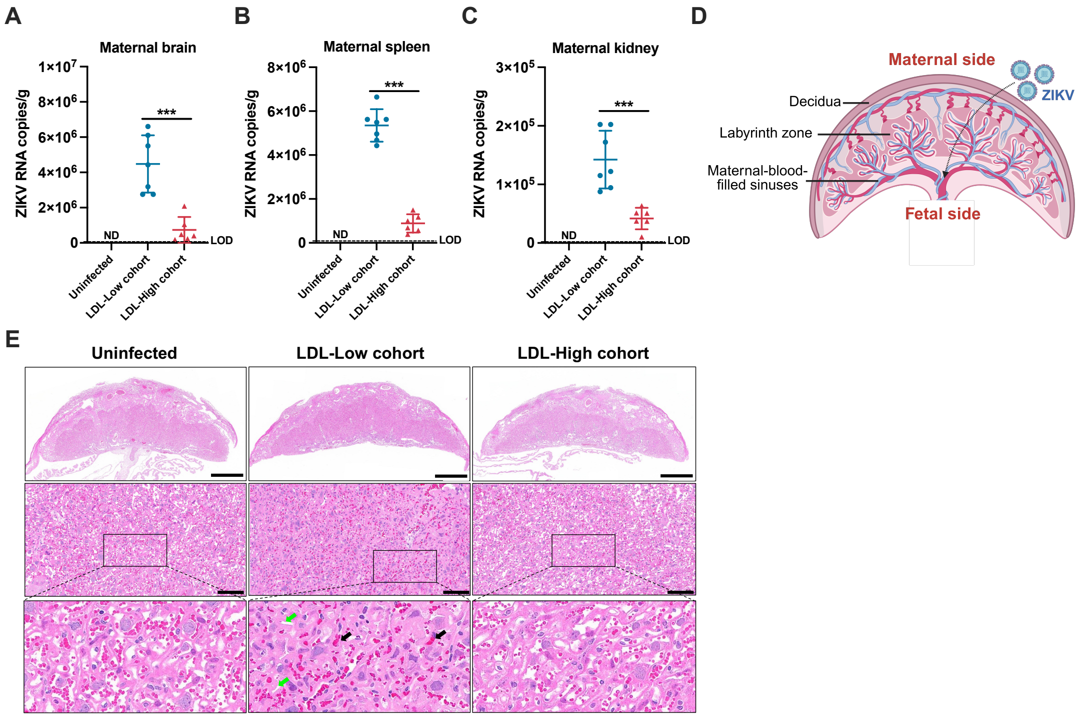


**Supplemental figure 8. Maternal ZIKV infection profiles and placenta histopathology.**

**A–C,** ZIKV RNA copies per gram of tissue (copies/g) in the maternal brain (**A**), spleen (**B**), and kidney (**C**) at E18.5, quantified by Q-RT-PCR (n = 6-7). **D**, Schematic diagram of the mouse placental structure. **E,** Representative H&E-stained placental sections. Black arrows indicate focal vascular congestion and hemorrhage within the placental labyrinth zone; green arrows indicate trophoblast degeneration and disorganization of the labyrinthine architecture. Data are mean ± s.d. from three independent experiments. Statistical analysis was performed using two-tailed unpaired Student's t-tests (**A–C**). $*P < 0.05$; $**P < 0.01$; $***P < 0.001$
